# Plasticity degeneracy underlies flexible formation and reconfiguration of spatial representations in hippocampal granule cells

**DOI:** 10.64898/2026.09.16.751981

**Authors:** Sanjna Kumari, Rishikesh Narayanan

**Affiliations:** Cellular Neurophysiology Laboratory, Molecular Biophysics Unit, Indian Institute of Science, Bengaluru 560012, India

**Keywords:** degeneracy, granule cells, hippocampus, heterogeneities, intrinsic plasticity, place cells, synaptic plasticity

## Abstract

Learning and memory require neural representations that are simultaneously robust and adaptable. How do neural systems reconcile these opposing demands of stability and flexibility? Here, we use hippocampal granule cells, which transform spatially diffuse entorhinal inputs into sparse sharply-tuned spatial representations, to address this fundamental question using emergence, stabilization, remapping, and suppression of place-fields as ideal systems to study stability-flexibility balance. Employing an unbiased population-based approach involving different strengths of plasticity in several components, we show that sharply tuned place-cell firing does not arise from a unique plasticity mechanism. We employed a population of heterogeneous, experimentally constrained biophysical models of dentate gyrus granule cells receiving cortical grid-field and contextual inputs to demonstrate that spatially restricted synaptic potentiation is insufficient to selectively route spatial information. Instead, each of the four distinct target transitions, namely reliable emergence, stabilization, remapping, and suppression of place-fields, were achievable by coordinated plasticity spanning excitatory synapses and several intrinsic ion channels. Across each of these four transitions, we identify numerous mechanistically distinct plasticity combinations that generate equivalent functional outcomes. These solutions are non-random and highly diverse, exhibiting weak dependence among individual plasticity components despite converging onto constrained functional states, through a combination of targeted amplification and global suppression of spatial firing. Our findings reveal extensive plasticity degeneracy in hippocampal spatial coding and suggest that flexible neural representations emerge not from unique plasticity rules involving one single component, but from a repertoire of alternative plasticity routes that are capable of implementing the same computation.

## INTRODUCTION

Neural representations must remain stable enough to support reliable memories while retaining the flexibility required to incorporate new experiences. This stability-flexibility dilemma is particularly evident in the hippocampus, where spatial representations of the external world can emerge, strengthen, weaken, remap, or disappear as animals learn and interact with changing environments. What cellular mechanisms mediate such adaptive yet robust representations in the brain? Hippocampal place cells provide a powerful system for addressing this question. Since their discovery (O’Keefe and Dostrovsky, 1971), place cells have served as a canonical model of memory-related neural representations, demonstrating how specific patterns of neuronal activity encode an animal’s position in space. The most attractive feature of place fields is that they are not static entities but are endowed with dynamic transitions that balance representational stability with ongoing plasticity. They have been shown to emerge *de novo* from silent cells, stabilize across repeated experiences, remap to new locations, or be selectively suppressed depending on specific requirements (Colgin et al., 2008; Moser et al., 2008; Hartley et al., 2014; Moser et al., 2015; Moser et al., 2017; Fenton, 2024).

The dentate gyrus (DG) plays a critical role in this process (Danielson et al., 2017; GoodSmith et al., 2017; Jung et al., 2019; Zhang et al., 2020; Cholvin et al., 2021; GoodSmith et al., 2022; Cholvin and Bartos, 2026). As the principal gateway of cortical information into the hippocampus, DG granule cells receive convergent inputs from the medial and lateral entorhinal cortices, which convey complementary grid-field and contextual information (Hafting et al., 2005; Hargreaves et al., 2005; Andersen et al., 2006; Amaral et al., 2007; Deshmukh et al., 2010; Deshmukh and Knierim, 2011; Yoganarasimha et al., 2011; Tsao et al., 2013; Yoon et al., 2016; Wang et al., 2018). How do granule cells selectively route information from such diverse and spatially diffuse inputs to generate sharply localized single-field firing (Danielson et al., 2017; GoodSmith et al., 2017; Senzai and Buzsaki, 2017; Zhang et al., 2020; Cholvin et al., 2021; GoodSmith et al., 2022; Cholvin and Bartos, 2026)? A prevailing view is that place-field formation and reconfiguration are driven primarily by location-dependent plasticity at active excitatory synapses. However, given the unique set of afferent inputs to the DG where afferent inputs are spatially diffuse and drive responses at multiple locations, targeted location-dependent synaptic plasticity would also result in off-field responses.

An additional layer of complexity arises from the extensive heterogeneity that characterizes hippocampal circuits. At the input end, entorhinal inputs vary in their spatial and contextual tuning (Hafting et al., 2005; Hargreaves et al., 2005; Deshmukh et al., 2010; Deshmukh and Knierim, 2011; Yoganarasimha et al., 2011; Tsao et al., 2013; Yoon et al., 2016; Wang et al., 2018). In addition, granule cells themselves differ substantially in intrinsic physiological properties and ion-channel expression (Beining et al., 2017; Mishra and Narayanan, 2019, 2020; Kumari and Narayanan, 2024, 2026). As neuronal output is determined not merely by synaptic strength but also by intrinsic conductances that regulate dendritic integration, excitability, and spike generation (Schmidt-Hieber et al., 2007; Krueppel et al., 2011; Lee et al., 2012; Hsu et al., 2018; Zhang et al., 2020), this heterogeneity is critical in driving selective routing of spatial information. Importantly, behaviorally relevant patterns of activity are known to induce conjunctive changes in both synaptic and intrinsic properties in DG granule cells (Greenstein et al., 1988; Pavlides et al., 1988; Shors and Dryver, 1994; Beck et al., 2000; Davis et al., 2004; McHugh et al., 2007; Larson and Munkacsy, 2015; Mishra and Narayanan, 2022), suggesting that spatial representations may emerge through interactions among multiple forms of plasticity. The question of whether such coordinated plasticity recruiting distinct molecular mechanisms could result in the emergence or reconfiguration of place fields remains unresolved.

We hypothesized that sharply tuned place cells flexibly emerge and reconfigure through conjunctive mechanisms of synaptic and intrinsic plasticity that couple targeted *local enhancement* of firing at specific spatial locations with *global reduction* in excitability to suppress non-specific firing. We tested this hypothesis using morphologically and biophysically detailed models of hippocampal granule cells that were extensively constrained by experimental electrophysiological measurements. We examined how heterogeneous spatial and contextual inputs are transformed into sharply tuned place-cell responses and whether distinct forms of place-field plasticity require unique or multiple mechanistic solutions. We investigated four fundamental target transitions in spatial coding: emergence of a new place field, stabilization of an existing field, remapping to a new location, and suppression of a spurious field. We stochastically searched a physiologically relevant plasticity space by testing several thousands of conjunctive plasticity combinations that could yield each of these fundamental target transitions.

We demonstrate that targeted synaptic potentiation alone is insufficient to reliably generate sharply tuned single-field responses. Instead, successful routing of spatial information requires coordinated plasticity involving both excitatory synapses and intrinsic ion-channel conductances. Across all target transitions, we identify multiple non-unique and non-random combinations of synaptic and intrinsic plasticity that yielded the specific target transition, revealing extensive plasticity degeneracy in the emergence and reconfiguration of hippocampal spatial representations. These observations identify plasticity degeneracy as a foundational organizing principle in the flexible formation and stable reconfiguration of neural representations.

## METHODS

### Heterogeneous model population of dentate gyrus granule cells

For all simulations reported in this study, we used the base model and the entire heterogeneous population of 141 models of dentate gyrus (DG) granule cells (GC) that were validated against 17 characteristic electrophysiological measurements in an earlier study (Kumari and Narayanan, 2024). The details of model specifications are identical to the earlier study (Kumari and Narayanan, 2024), which were derived originally from (Beining et al., 2017), with key details reproduced below. The GC morphology was stratified into seven sections: outer molecular layer (OML), middle molecular layer (MML), inner molecular layer (IML), granule cell layer (GCL), soma, axon initial segment (AIS), and axon. The passive and active properties were variable across sections (Supplementary Table S1). Spines were accounted for by scaling the specific membrane resistance and specific membrane capacitance in the IML by a factor of 1.45, and in the MML and OML by a factor of 1.9 (Beining et al., 2017). Leak channels were distributed non-homogeneously across sections (Hervieu et al., 2001; Gabriel et al., 2002; Yarishkin et al., 2014; Beining et al., 2017) by altering the leak conductance *g*_pas_ (Supplementary Table S1).

We incorporated 15 active conductances into the model (Kumari and Narayanan, 2024), with their gating kinetics and subcellular distributions adapted and re-tuned from the original model (Beining et al., 2017) (Supplementary Table S1). An inward-rectifier potassium channel (*K*_ir_) (Lopatin et al., 1995; Yan and Ishihara, 2005; Panama and Lopatin, 2006; Ishihara and Yan, 2007) was set to be present in all sections, with location-dependent distribution of conductance. An 8-state sodium channel (Na) with region-dependent densities was incorporated into the model with the highest density in the AIS and lower densities in dendrites and soma (Kress et al., 2010; Schmidt-Hieber and Bischofberger, 2010). Inactivating voltage-gated potassium channels K_v_1.1 (Christie et al., 1989) and K_v_1.4 (Wissmann et al., 2003) were present in AIS and axon, whereas K_v_4.2 (Barghaan et al., 2008) was localized to dendrites. The delayed-rectifier K_v_3.4 channels (Rudy et al., 1991; Schroter et al., 1991; Rettig et al., 1992; Vega-Saenz de Miera et al., 1992; Riazanski et al., 2001) were incorporated only into the axonal and AIS compartments. *M*-type potassium channels (K_v_7.2/7.3) (Mateos-Aparicio et al., 2014) were localized to the axon and the AIS. The hyperpolarization-activated cyclic nucleotide gated (HCN) non-specific cationic channels (Stegen et al., 2012) were inserted in the dendrites. Voltage-gated *N*-type (Ca_v_2.2) (Fox et al., 1987) and *T*-type calcium channels (Ca_v_3.2) (Burgess et al., 2002) were distributed across all compartments. Voltage-gated *L*-type calcium channels (Ca_v_1.2/1.3) (Evans et al., 2013) were inserted such that Ca_v_1.3 was present in all compartments and Ca_v_1.2 spread across all sections except for the axon. The calcium-dependent big-conductance potassium (BK) channels (Jaffe et al., 2011) were inserted into the soma and axon. The calcium-dependent small-conductance potassium (SK) channels (Hirschberg et al., 1999; Solinas et al., 2007) were incorporated in all the sections. We specifically note the differential localization profiles of Kir (global localization), Na (predominantly axonal localization at the AIS), and HCN (dendritic localization) channels, as plasticity profiles in these conductances will later be defined by their localization.

A majority of these ion channels were modeled using Hodgkin-Huxley dynamics (Hodgkin and Huxley, 1952). *A*-type K_v_4.2 potassium channels followed a 15-state Markovian model. *T*-type Ca_v_3.2 calcium and sodium channels were modeled as 8-state Markovian models. SK and K_ir_ channels were both modeled as 6-state Markov models. Calcium buffer shell model was modified from (Anwar et al., 2014). The Ca^2+^ decay time constant was set to 43 ms (Jackson and Redman, 2003) in the axon and 240 ms (Stocca et al., 2008) in all other compartments. Sodium reversal (*E*_Na_) was set to 50 mV and potassium reversal (*E*_K_) to –80 mV. Sodium channels (Na8st) were introduced in the dendritic compartments in GCL, IML, MML and OML strata to accommodate active dendrites in GCs (Krueppel et al., 2011). All model parameters and their respective base values are listed section wise in Supplementary Table S1.

Models were compartmentalized using the *d_λ_* rule (Carnevale and Hines, 2006), whereby each compartment in the model was set to be less than 10% of the space constant of the neuronal section, computed at 100 Hz. This compartmentalization process yielded a total of 233 compartments in the base model, of which 163 were somato-dendritic compartments. In addition to the base model, we also used the entire heterogeneous population of 141 valid GC models that were generated using stochastic search process (Kumari and Narayanan, 2024).

### Electrophysiological measurements

The base model and the 141 heterogeneous models were validated against 17 electrophysiological signature measurements (Supplementary Tables 2–3) reported for dorsal and intermediate granule cells (Mishra and Narayanan, 2020; Kumari and Narayanan, 2026). The measurements were computed using well-established procedures described below and are identical to corresponding electrophysiological quantifications (Mishra and Narayanan, 2020; Kumari and Narayanan, 2024, 2026).

### Sub-threshold intrinsic measurements

Resting membrane potential (*V_RMP_*) was measured as the potential at which the membrane rested when no current is injected. *V_RMP_* was calculated as the mean of recorded voltage for the last 50 ms of a 1-s simulation performed in the absence of current injection. All sub- and supra-threshold measurements were performed after an initial delay of 1 s to allow *V_RMP_* to reach steady-state value. Input resistance (*R_in_*) was measured as the slope of a linear fit to the steady-state *V–I* plot obtained by injecting sub-threshold current pulses of amplitudes spanning –50 to +50 pA, in steps of 10 pA. Percentage sag was measured from the voltage response of the cell to a hyperpolarizing current pulse of –100 pA for 1000 ms and was defined as: 100[1 − (*V_SS_* / *V_peak_*)], where *V_SS_* and *V_peak_* depicted the steady-state and peak voltage deflection from *V_RMP_*, respectively. To assess temporal summation, five α-excitatory postsynaptic currents (α-EPSCs) with 50 ms intervals were injected into the somatic compartment. Temporal summation ratio (*S_α_*) was computed as *E_last_* /*E_first_*, where *E_last_* and *E_first_* are the amplitudes of last and first α-excitatory postsynaptic potentials, respectively, recorded in response to the injection of 5 α-EPSCs. The chirp stimulus used for characterizing the impedance profiles was a sinusoidal current of constant amplitude below firing threshold, with its frequency linearly spanning 0–15 Hz in 15 s. The magnitude of the ratio of the Fourier transform of the voltage response to the Fourier transform of the Chirp stimulus yielded the impedance amplitude profile (Narayanan and Johnston, 2008):

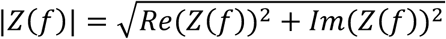

where *Re*(*Z*(*f*)) and *Im*(*Z*(*f*)) were the real and imaginary parts, respectively, of the impedance *Z* as a function of frequency, *f*. The peak value of impedance across all frequencies was measured as the maximum impedance amplitude |*Z*|_max_. The frequency at which the impedance amplitude reached its maximum value was defined as the resonance frequency (*f_R_*). Resonance strength (*Q*) was measured as the ratio of the maximum impedance amplitude to the impedance amplitude at 0.5 Hz. Impedance phase *φ*(*f*) was computed as:

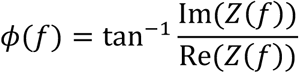

Total inductive phase, Φ_L_, defined as the area under the inductive part of *φ*(*f*) (Fig. 1*G*) was defined as (Narayanan and Johnston, 2008):

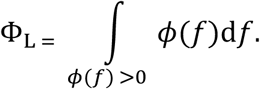

**Figure 1:**
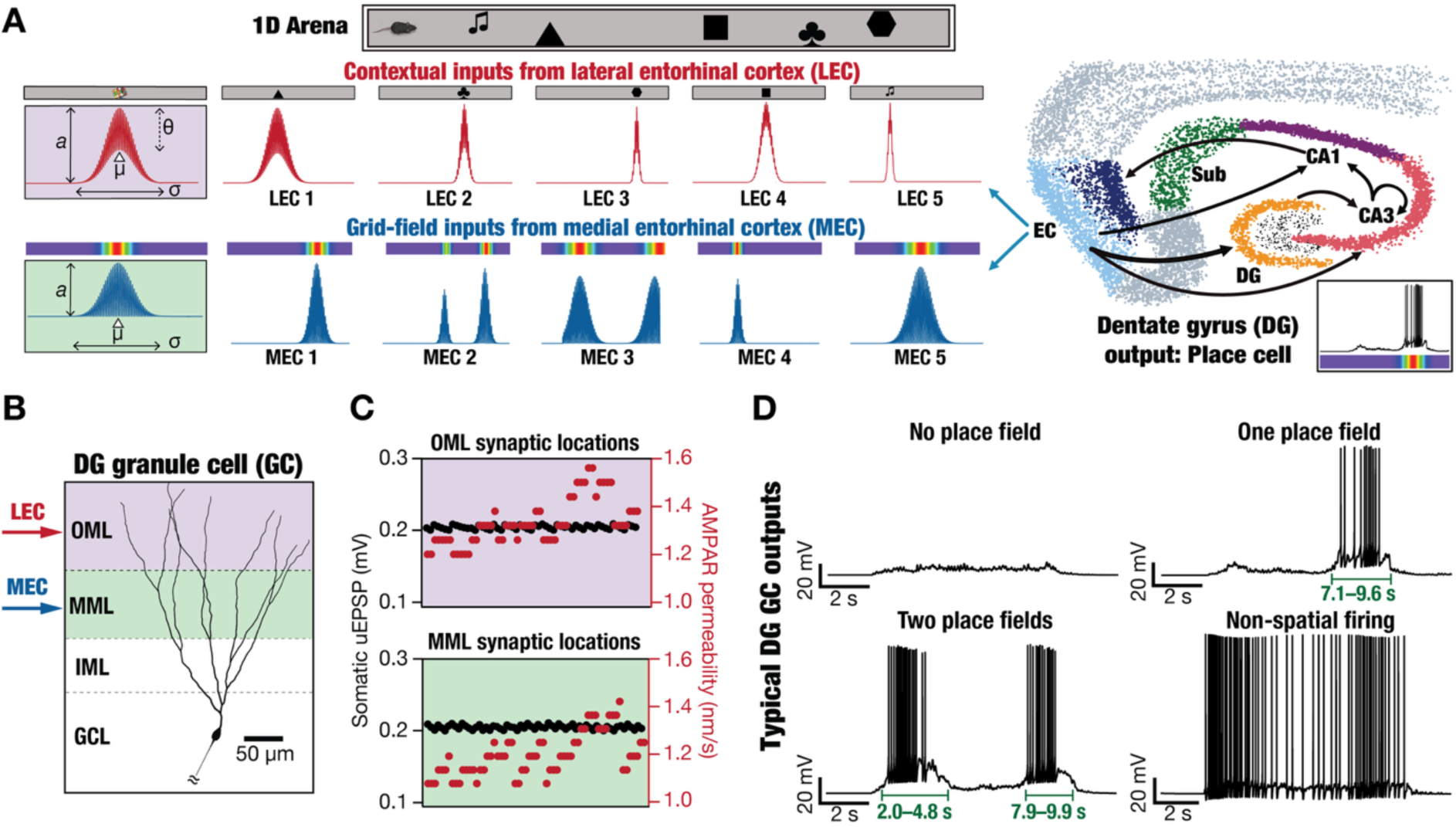
Typical firing responses of granule cell models to conjunctive distally innervating contextual LEC inputs and proximally innervating spatial MEC inputs. (A) Granule cells of the dentate gyrus receive contextual inputs from the lateral entorhinal cortex, LEC (represented as firing dependent on object-location along the 1D arena) and spatially selective grid-field inputs from the medial entorhinal cortex, MEC. The firing profile of LEC inputs (*Left, Top*) was represented by four parameters defining the amplitude of Gaussian-modulated firing (*a*), standard deviation of Gaussian (*σ*), the mean of the Gaussian (*μ*) and the degree of theta modulation (*θ*). The grid-field firing of MEC inputs (*Left, Bottom*) was represented with three parameters, amplitude of gaussian (*a*), standard deviation of Gaussian (*σ*) and the mean of the Gaussian (*μ*). Five examples of typical LEC and MEC inputs are shown. Note that there can be more than one grid-field for specific cells with different parametric values. *Right bottom*, typical granule cell firing showing a single place field. This study delineates the mechanisms behind the emergence of such single place-field firing in DG granule cells, given the diversity of inputs that impinge on the DG granule cell. (B) Morphologically realistic conductance-based model of DG GC with different layers labelled. The model receives 25 randomly generated LEC-like inputs and MEC-like inputs (as shown in panel A), each onto dendritic locations within the outer and middle molecular layers respectively. (C) AMPAR permeability values (*P*_AMPAR_, right axis) at each synaptic location along the outer molecular layer (*Top*, OML) and middle molecular layer (*Bottom*, MML) were set such that the somatic unitary excitatory postsynaptic potentials (uEPSPs, left axis) were around 0.2 mV, irrespective of synaptic location. (D) Voltage traces showing the different types of granule cell outputs generated with disparate sets of LEC-MEC input combinations impinging on the dendritic tree. Different sets of inputs yielded outputs with no place field, one place field, two place fields, or non-spatial firing. The durations marked at the bottom of the “one place field” and “two place fields” traces represent the specific locations where firing was high.

### Supra-threshold intrinsic measurements

Supra-threshold measurements were obtained through depolarizing current injections, with amplitudes large enough to elicit action potentials (AP), into the cell resting at *V_RMP_* (Mishra and Narayanan, 2019, 2020; Kumari and Narayanan, 2026). AP firing frequency was computed by counting the number of spikes obtained during a 1000 ms current injection. Current amplitude of these pulse-current injections was varied from 0 pA to 250 pA in steps of 50 pA, to construct the firing frequency *vs*. injected current (*f* − *I*) plot. Various AP related measurements were derived from the voltage response of the cell to a 250-pA pulse-current injection. The temporal distance between the timing of the first spike and the time of current injection was defined as latency to first spike (*T*_1AP_). The duration between the first and the second spikes was defined as the first inter-spike interval (*T*_1ISI_). AP amplitude (*V_AP_*) was computed as the difference between the peak voltage of the first spike and *V_RMP_*. AP half-width (*T*_APHW_) was the temporal width measured at the half-maximal points of the AP peak with reference to *V_RMP_*. The maximum (*dV*/*dt*|*_max_*) and minimum (*dV*/*dt*|*_min_*) values were calculated from the temporal derivative of the first action potential obtained with 250-pA current injection. The voltage in the AP trace corresponding to the time point at which the *dV*/*dt* crossed 20 V/s was defined as AP threshold (*V_t_*_ℎ_). The sub- and supra-threshold measurements of the base model and their respective experimentally derived bounds are listed in Supplementary Tables S2–S3. It may be noted that all base model measurements are within their respective electrophysiological bounds.

### Synapse model

Excitatory synapses from the medial and lateral entorhinal cortices impinged on granule cell dendrites within the middle molecular layer (MML) and the outer molecular layer (OML) of the DG, respectively (Fig. 1*A–B*). These synapses were modeled with co-localized AMPA-NMDA receptors (Ye et al., 2005; Krueppel et al., 2011) and were randomly distributed across dendritic compartments within the MML and OML. The default value of NMDAR:AMPAR ratio was set to 1. The ionic current through these receptors were modeled using the Goldman– Hodgkin–Katz (GHK) convention (Goldman, 1943; Hodgkin and Katz, 1949; Narayanan and Johnston, 2010). The current through NMDA receptors reflected their voltage-dependence and ionic composition, carried by sodium, potassium, and calcium ions:

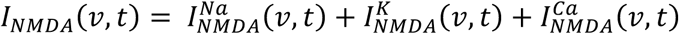

where,

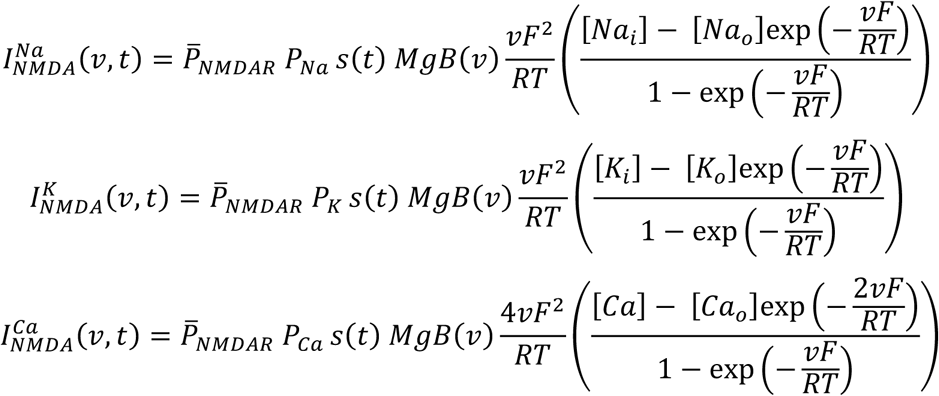

where *P̄_NMDAR_* defined the maximum permeability of the NMDA receptor. The relative permeability values for Na+ and K+ were set as *P*_Na_ = *P*_K_ = 1 and for calcium, *P*_Ca_ = 10.6. The intra- and extra-cellular concentrations for the different ions were set as: [*Na*]*_i_* = 18 mM, [*Na*]*_o_* = 140 mM, [*K*]*_i_* = 140 mM, [*K*]*_o_* = 5 mM, [*Ca*]*_i_* = 100 nM and [*Ca*]*_o_* = 2 mM. These ionic concentrations ensured that the reversal potentials for both AMPA and NMDA were set at 0 mV. The role of magnesium in regulating the activity of the NMDAR was accounted by the *MgB*(*v*) factor (Jahr and Stevens, 1990):

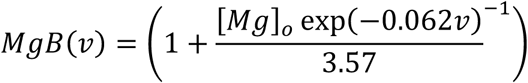

where [*Mg*]_o_ = 2 mM. *s*(*t*) governed the kinetics of the NMDA receptor as follows (Jahr and Stevens, 1990):

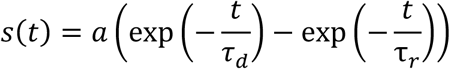

where *a* defined a normalization factor that ensured that 0 ≤ *s*(*t*) ≤ 1. τ*_r_* (= 7 ms) and *τ_d_* (= 275 ms) represented the rise time and decay time constant, respectively (Ye et al., 2005; Krueppel et al., 2011). The AMPAR current was modeled following the GHK convention, and was driven by sodium and potassium:

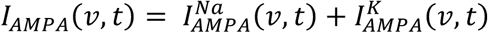

where

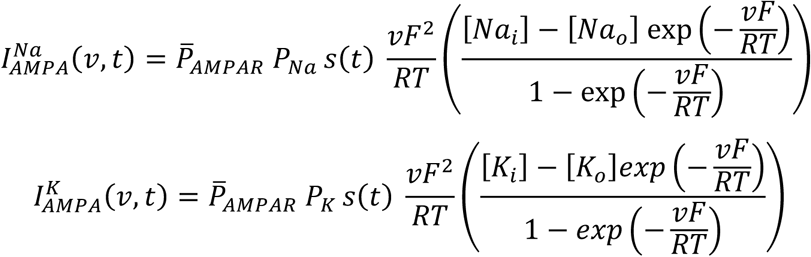

where, *P̄_AMPAR_* is the maximum permeability of the AMPA receptors, with *P*_Na_ = *P*_K_ = 1. The rise and decay time constants of AMPAR were τ*_r_* (= 2 ms) and *τ_d_* (= 10 ms) (Ye et al., 2005). The AMPA permeabilities were tuned for each dendritic location along the MML and OML to ensure that the unitary somatic EPSP amplitude was set at 0.2 mV (Krueppel et al., 2011), irrespective of dendritic location of the synapse (Fig. 1*C*).

### Afferent inputs to the granule cell dendrites

A probability density function modeled as summations of one or two Gaussian-modulated cosinusoids governed the spikes arriving onto synapses from the entorhinal cortices onto the dendritic tree. The cosinusoidal frequency, set at 8 Hz, mimicked the theta activity present in hippocampal formation during exploratory behavior whereas the Gaussian mimicked the transient increase in excitatory drive owing to context (for LEC inputs) or the animal entering the cell’s grid field. The formulation for inputs from LEC is similar to those used for modeling place-field inputs to CA1 pyramidal neurons (Basak and Narayanan, 2018, 2020; Roy and Narayanan, 2021), but accounting for multiple objects or multiple grid fields in a single traversal as well as for the differential theta modulation of firing in LEC *vs*. MEC cells (Hafting et al., 2005; Hargreaves et al., 2005; Deshmukh et al., 2010; Deshmukh and Knierim, 2011; Yoganarasimha et al., 2011; Tsao et al., 2013; Yoon et al., 2016; Wang et al., 2018). The velocity of the animal traversing the arena was assumed to be constant, implying that space and time are interchangeably used. We use time as the default independent variable, with the 1D arena designed to be spanned over a period of 10 seconds.

### Inputs from the lateral entorhinal cortex

The contextual lateral entorhinal cortex inputs (Deshmukh and Knierim, 2011; Tsao et al., 2013; Wang et al., 2018) were modeled using a sum of two Gaussian-modulated cosinusoids. Each synapse from LEC cells onto the granule cell received presynaptic spikes with the probability of occurrence at time *t* defined by:

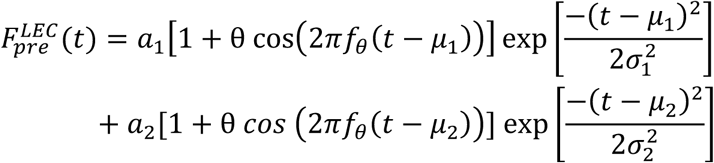

where, *a*_1_ and *a*_2_ (range: 0.1–10 Hz) are the maximum presynaptic firing rates, *μ*_1_ (range: 2– 6 s) and *μ*_2_ (to avoid strongly overlapping fields, *μ*_2_ = *μ*_1_ + *σ*_1_ + *σ*_2_ + *K*, where *K* was randomly picked from the range 0–4 s) are the center location of spatially localized firing, *σ*_1_and *σ*_2_ (range: 0.1–1 s) are the widths of the first and second firing fields. *f_θ_* = 8 Hz defined theta-frequency modulation of LEC-cell firing rate. *θ* (range: 0.1–0.5) determined the degree of theta modulation of the LEC inputs, which has been shown to be less than theta modulation of MEC firing (Deshmukh et al., 2010). All parameters were randomly sampled from independent uniform distributions within their respective ranges, for 25 distinct LEC inputs. The probability of an LEC afferent input eliciting action potentials for two objects (*i.e.*, two firing fields) within the linear arena was set at 20%. Therefore, for 80% of the cases where there was only one firing field, *a*_2_ was set to zero. Examples of LEC inputs generated are shown in Fig. 1*A*. The 25 distinct inputs from afferent LEC neurons impinged on synapses that were randomly distributed within the outer molecular layer (OML) of the granule-cell dendritic tree.

### Inputs from the medial entorhinal cortex

The grid-like medial entorhinal cortex (MEC) inputs (Hafting et al., 2005; Yoganarasimha et al., 2011; Yoon et al., 2016) were modeled using a sum of two Gaussian-modulated cosinusoids. Each synapse from MEC cells onto the granule cells received presynaptic spikes with the probability of occurrence at time *t* defined by:

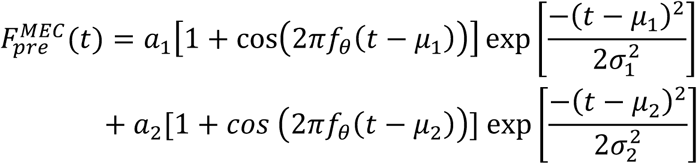

where, *a*_1_ and *a*_2_ (range: 0.1–10 Hz) are the maximum presynaptic firing rates, *μ*_1_ (range: 2– 6 s) and *μ*_2_ (to avoid strongly overlapping fields, *μ*_2_ = *μ*_1_ + *σ*_1_ + *σ*_2_ + *K*, where *K* was randomly picked from the range 0–4 s) are the center location of spatially localized firing, *σ*_1_ and *σ*_2_ (range: 0.1–1 s) are the widths of the first and second firing fields. *f_θ_* = 8 Hz defined theta-frequency modulation of MEC-cell firing rate. The theta modulation was 100% in this case. All parameters were randomly sampled from independent uniform distributions within their respective ranges, for 25 distinct MEC inputs. The probability of an MEC afferent input eliciting action potentials at two grid-fields within the linear arena was set at 20%. Therefore, for 80% of the cases where there was only one firing field, *a*_2_ was set to zero. Examples of MEC inputs generated are shown in Fig 1*A*. The 25 distinct inputs from afferent MEC neurons impinged on synapses that were randomly distributed within the middle molecular layer (MML) of the granule-cell dendritic tree. The synaptic strengths associated with these inputs were defined by the AMPAR permeability value for the location where the synapse impinged (Fig. 1*C*), with the NMDAR permeability for the synapse defined by the NMDAR:AMPAR ratio.

### Measurements to quantify granule-cell spikes and their spatial localization

The postsynaptic responses to fifty synapses receiving heterogeneous afferent inputs from the lateral and medial entorhinal cortices were recorded as somatic voltages (Fig. 1*D*). Smooth instantaneous firing rate profiles were obtained from action potential timings using convolution with a Gaussian kernel (*σ* = 200 ms). Several measurements were derived from this instantaneous firing rate profile: (1) total firing rate across the arena, *f*_arena_, computed as the ratio between the total number of action potentials across the entire arena and the simulation time (10 seconds). (2) place-field firing rate, *f*_PF_, computed as the ratio between the total number of action potentials within a designated place field and the duration of the place field. To determine the extent of a place field, we identified the maximum of the instantaneous firing rate across the arena and then located the right and left cutoff points where the firing rate dropped to zero. These cutoff points were extended by 100 ms on either side to obtain the place-field bounds. (3) spatial selectivity coefficient, *C_SS_*, defined as the ratio between the peak in-field (within the place field) and peak out-of-field (rest of the arena) firing rates (Basak and Narayanan, 2018). If the peak in-field firing rate was zero, *C_SS_* was set to zero and if the peak out-of-field firing rate was zero, *C_SS_* was set to 10²⁰, a large value so that these scenarios can be plotted. Sharply tuned place-cell responses are characterized by a large value of *C_SS_*.

#### Multi-parametric multi-objective stochastic search (MPMOSS) on the plasticity space

We employed a multi-parametric multi-objective stochastic search (MPMOSS) algorithm to identify plasticity combinations that will result in the emergence of sharply tuned single place-field firing. The multiple parameters in this scenario were fold increases in each of the four plasticity components mentioned above (Δ*P_AMPAR_*, Δ*g_Na_*, Δ*g*_HCN_, and Δ*g_Kir_*). For each iteration, a random sample of each of these components was generated from their respective uniform distributions (Table 2). The model was subjected to this combination of random plasticity based on the targeted location of the synapse (Table 1) as well as the three sets of ionic conductances at their specific loci of plasticity (Table 2). The firing responses of the model across the arena were recorded to evaluate *f_PF_* and *C_SS_*. The plasticity combination was declared to be a valid place-cell model if the arena firing response satisfied the bounds specified for both *f_PF_* and *C_SS_* for the specific target transition required (Table 3). All sub- and supra-threshold intrinsic measurements were computed for all valid place-cell models, for each of the four target transitions. The parameters and intrinsic measurements of these valid place-cell models were further analyzed using Pearson’s correlation and dimensionality reduction techniques. Of the 17 total measurements that we made (Supplementary Tables S2–S3), we ignored two measurements (total inductive phase and firing rate for 50 pA) for dimensionality reduction analyses, as both of them were close to zero in most models.

**Table 1:** Target place-field locations for the four types of target transitions (see Fig. 1*D*).

|  | Target transition | Location of existing place field(s) | Location of target place field |
| --- | --- | --- | --- |
| 1. | Convert silent cell to place cell | — | 3–5 s |
| 2. | Stabilize existing place field | 7.1–9.6 s | 7.1–9.6 s |
| 3. | Spatially remap place field | 7.1–9.6 s | 3–5 s |
| 4. | Suppress second place field | 2–4.8 s & 7.9–9.9 s | 2–4.8 s |

**Table 2:**
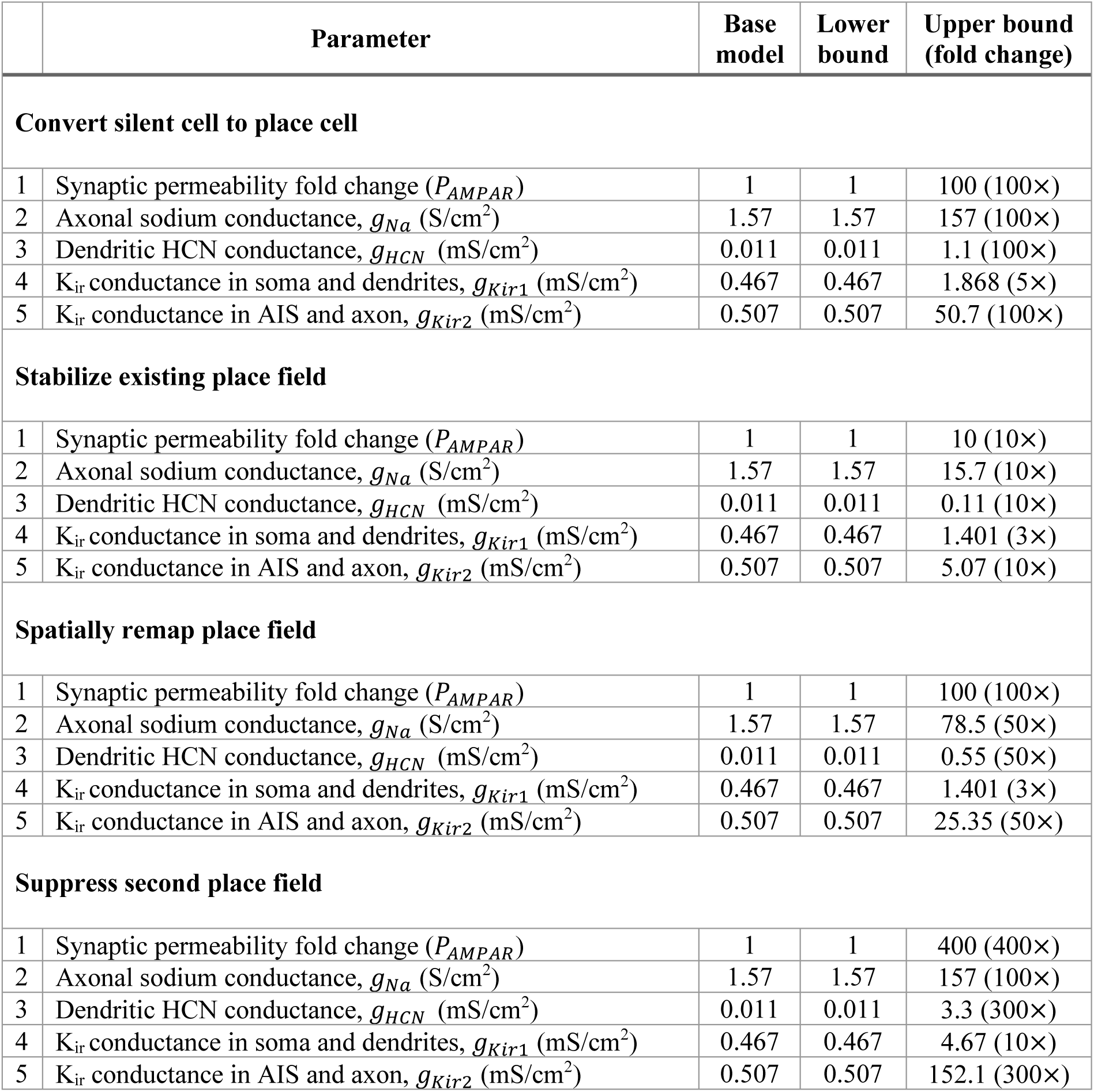
Bounds on plasticity parameters for MPMOSS to generate random plasticity combinations, for each of the four target transitions.

**Table 3:** Validation criteria for defining the four types of targeted transitions using MPMOSS.

|  | Types of responses | Threshold |
| --- | --- | --- |
| **Convert silent cell to place cell** |  |  |
| 1. | Spatial selectivity coefficient, $C_{SS}$ | $> 10$ |
| 2. | In-field firing rate, $f_{PF}$ | $> 3$ Hz |
| **Stabilize existing place field** |  |  |
| 1. | Spatial selectivity coefficient, $C_{SS}$ | $> 10$ |
| 2. | In-field firing rate, $f_{PF}$ | $> 6$ Hz |
| **Spatially remap place field** |  |  |
| 1. | Spatial selectivity coefficient, $C_{SS}$ | $> 10$ |
| 2. | In-field firing rate, $f_{PF}$ | $> 6$ Hz |
| **Suppress second place field** |  |  |
| 1. | Spatial selectivity coefficient, $C_{SS}$ | $> 10$ |
| 2. | In-field firing rate, $f_{PF}$ | $> 3$ Hz |

### Computational Details

All simulations were performed using the NEURON programming environment (Hines and Carnevale, 1997) at 34° C. The simulation step size was set as 25 μs. Data analysis and plotting of graphs were done using custom-built software written in MATLAB or Igor Pro programming environment (WaveMetrics Inc., USA). To avoid false interpretations and to emphasize heterogeneities in simulation outcomes, the entire range of measurements are reported in figures rather than providing only summary statistics (Marder and Taylor, 2011; Rathour and Narayanan, 2019).

## RESULTS

What mechanisms underlie the transformation of the diversity of spatial and contextual inputs from the entorhinal cortex to sharply tuned single-field firing outputs of hippocampal granule cells (Fig. 1*A*)? How do granule cells achieve such selective routing of spatial information, especially when afferent inputs from the entorhinal cortex elicit firing at several diverse locations and lack sharply tuned spatial selectivity at single spatial locations (Fig. 1*A*)? In addressing these questions, we employed a morphologically and biophysically detailed model population of hippocampal granule cells (Kumari and Narayanan, 2024) with ion-channel distributions explicitly constrained across distinct anatomical compartments (Supplementary Table S1) to individually match experimentally observed expression profiles (Beining et al., 2017; Kumari and Narayanan, 2024). This model population consisted of a base model as well as the entire heterogeneous population of 141 valid GC models (Kumari and Narayanan, 2024) generated through an unbiased stochastic search across a 45-parameter space (Supplementary Table S1). These models were generated after validation using 17 different intrinsic physiological measurements (Supplementary Table S2–S3), which matched with respective measurements from dorsal and intermediate hippocampal granule cells (Mishra and Narayanan, 2020; Kumari and Narayanan, 2026).

We modeled grid-like and contextual inputs from the medial (MEC) and lateral (LEC) entorhinal cortices, respectively, explicitly accounting for their firing profiles in 1D arenas and the extent of theta modulation of their firing rates (Hafting et al., 2005; Hargreaves et al., 2005; Deshmukh et al., 2010; Deshmukh and Knierim, 2011; Yoganarasimha et al., 2011; Tsao et al., 2013; Yoon et al., 2016; Wang et al., 2018). Importantly, MEC and LEC inputs were stochastically generated, encoding diverse spatial and contextual information along a linear track, rather than being preconfigured to fall into specific pre-defined locations (Fig. 1*A*). Afferent inputs from the medial and lateral entorhinal cortex (25 diverse inputs each) made contact through glutamatergic synapses located within the middle molecular layer (MML) and the outer molecular layer (OML), respectively, of intrinsically validated granule cell models (Fig. 1*B*). Synapses contained AMPA and NMDA receptors postsynaptically, with AMPA receptor permeability (density) set such that the somatic unitary EPSP amplitude was 0.2 mV (Krueppel et al., 2011), irrespective of the location of synaptic origin (Fig. 1*C*). Thus, our well-constrained models (Fig. 1*A–C*; Supplementary Tables S1–S3) explicitly accounted for the morphology and channel distributions of granule cells (Beining et al., 2017; Kumari and Narayanan, 2024), for several intrinsic physiological measurements that matched with dorsal granule cells especially (Kumari and Narayanan, 2024, 2026), for firing characteristics of afferent entorhinal inputs and their postsynaptic dendritic locations (Hafting et al., 2005; Hargreaves et al., 2005; Amaral et al., 2007; Deshmukh et al., 2010; Deshmukh and Knierim, 2011; Yoganarasimha et al., 2011; Tsao et al., 2013; Yoon et al., 2016; Wang et al., 2018), and unitary amplitudes of synapses impinging at different locations (Krueppel et al., 2011).

We recorded somatic voltage responses to disparate random sets of entorhinal inputs (each set contained 25 LEC and 25 MEC inputs, all of which were randomly generated) impinging on granule cell dendrites. Dendritic computation involving spatiotemporal integration of all afferent inputs eventually translated to somatic voltage deflections in the model neuron. Somatic voltage recordings from neurons receiving different sets of inputs revealed a diverse spectrum of naturally emergent firing patterns, which fell into one of four major categories (Fig. 1*D*): (a) silent cells that did not elicit action potentials at any location across the arena; (b) neurons that yielded a well-defined single field at a specific location within the arena; (c) neurons that manifested action potential firing at two well-delineated place field locations within the arena; and (d) neurons that showed global firing with no discernible pattern. These firing patterns are naturally emergent simply as a consequence of postsynaptic integration of afferent inputs through synapses that were normalized to yield similar unitary responses at the soma (Fig. 1*C*). These firing patterns and subthreshold responses showing the presence of spatial information across the arena at various baseline strengths are reminiscent of experimental observations demonstrating such patterns and responses in hippocampal place cells (Lee et al., 2012; Bittner et al., 2015; Bittner et al., 2017; Diamantaki et al., 2018; Magee and Grienberger, 2020; McKenzie et al., 2021; Milstein et al., 2021; Valero et al., 2022).

Of these four types of firing profiles, we ignored models that showed global firing with no discernible pattern. We considered the first three types of naturally emergent firing patterns and asked if these three types of firing patterns could be transformed into the physiologically observed firing pattern containing a single, sharply tuned place field within the arena.

### Target transitions required from different naturally emergent firing profiles to achieve sharply tuned, single-field place cells

Our target was to assess mechanisms that could transform a diversity of spatially non-selective entorhinal inputs onto a granule cell into sharply tuned, spatially localized, single-field firing. Based on the category of naturally emergent firing in response to the specific set of entorhinal inputs (Fig. 1*D*), this translated to four distinct transitions towards achieving our target (Fig. 2*A*):

1. **Convert silent cell to place cell**: The naturally emergent firing resulted in a silent cell. The target is to convert this into a single-field place cell, with action potential firing manifesting at an arbitrarily specified arena location.
2. **Stabilize existing place field**: The naturally emergent firing resulted in single-field place cell. The target is to stabilize this preexisting field at the same arena location, without introducing spurious action potential firing outside this field.
3. **Spatially remap place field**: The naturally emergent firing resulted in single-field place cell. The target is to spatially remap this place field to another arbitrarily specified arena location. Action potential firing within the preexisting field must be suppressed and sharply tuned single-field firing should emerge at the new location.
4. **Suppress second place field**: The naturally emergent firing resulted in action potential firing with two well-defined place fields. The target is to suppress one (arbitrarily chosen field) among the two while stabilizing the other. Action potential firing within one of the preexisting place fields must be suppressed and sharply tuned single-field firing should manifest at the other preexisting location.

**Figure 2:**
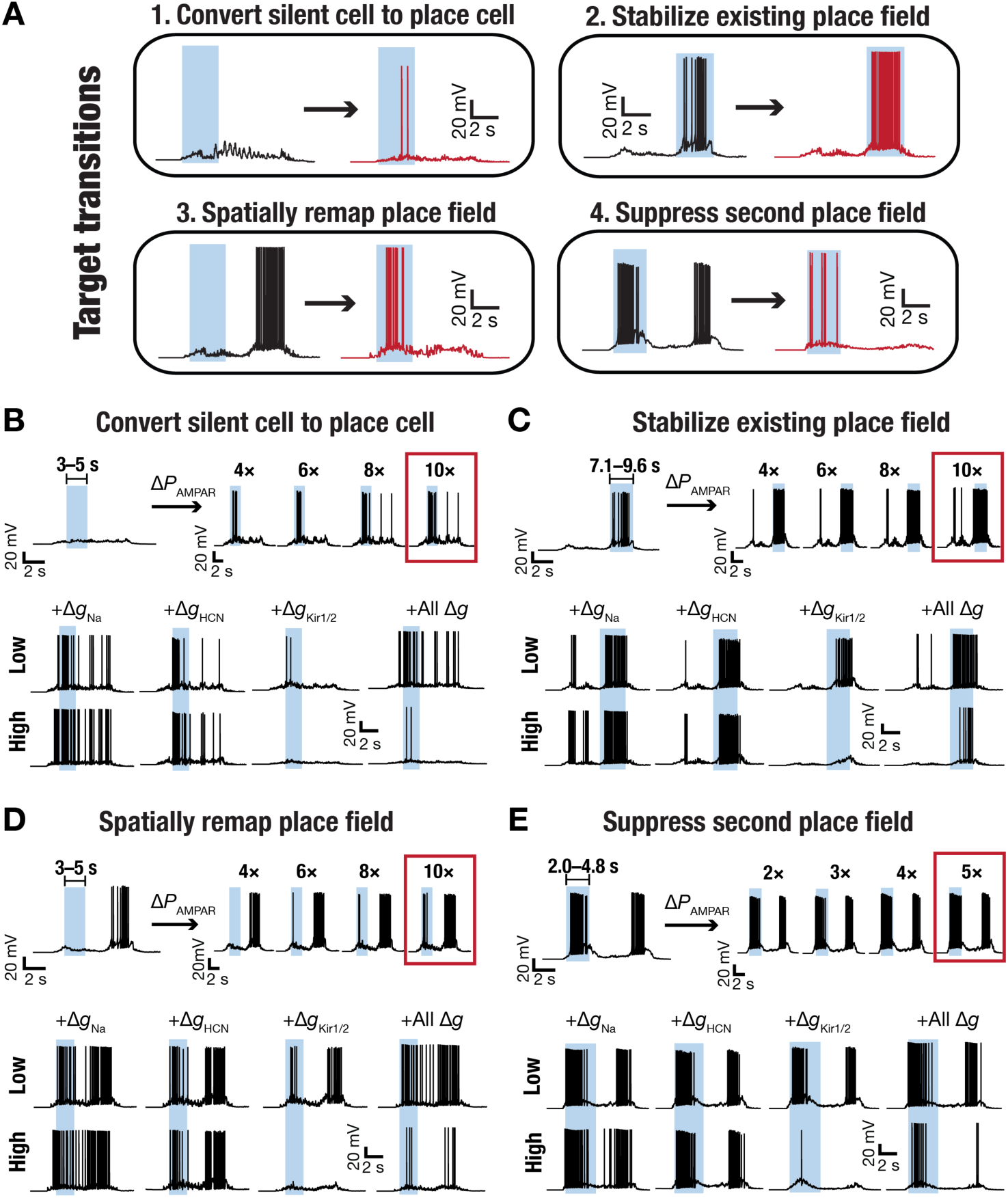
Diversity of the impact of concomitant changes in synaptic and intrinsic properties on the four target transitions. (A) The four target transitions, to be elicited through synaptic and/or intrinsic plasticity, from typical responses elicited by DG granule cells in response to EC inputs (shown in Fig. 1D). (B) **Convert silent cell to place cell**: *Top left*, Voltage response of a silent cell with no place fields along the 1D track. The shaded region with the bar on the top marks the target location where the place field should emerge. *Top Right*, Voltage responses of the same cell after increasing synaptic permeabilities associated with inputs aligned to the shaded region. Traces represent increase in AMPAR permeabilities (Δ*P_AMPAR_*) of those synapses by 4×, 6×, 8×, and 10× of their respective baseline values. *Bottom*, Voltage traces recorded when the model was subjected to 10× Δ*P_AMPAR_* (highlighted with a rectangle) was subjected to further intrinsic changes of: *Column 1*, 7× (Low) or 32× (High) increase in the sodium conductance (*g_Na_*) in the axonal initial segment; *Column 2*, 37× (Low) or 182× (High) increase in HCN conductance (*g_HCN_*) in dendrites; *Column 3*, 1.2× (Low) and 2× (High) inward rectifying potassium channel conductance values (*g_Kir_*) in soma and dendrites; and *Column 4,* 1.2× (Low) and 2× (High) increase in *g_Kir_* in soma and dendrites, but when *g_Na_* in AIS and *g_HCN_* in dendrites were set at higher value of 19× and 10×, respectively. (C–E) Same organization as Panel B for the three other target transitions: stabilize existing place field (C), spatially remap place field (D), and suppress second place field (E). Δ*P_AMPAR_* are different for each of the different targets and are marked above respective traces.

The end goal of all four target transitions is to achieve sharply tuned action potential firing within a single pre-specified spatial location, with the transition required to achieve that critically dependent on the naturally emergent firing. Therefore, the diversity of afferent input patterns onto individual granule cells and the consequent heterogeneity in naturally emergent (pre-transition) firing responses together constitute critical layers of diversity in building a sharply tuned, single-field place cell.

### Targeted synaptic plasticity of entorhinal synapses alone was insufficient to achieve sharply tuned single-field place fields

What plasticity mechanisms do we recruit to implement these transitions? Motivated by demonstrations of synaptic potentiation limited to synapses that were active during specific animal locations (Bittner et al., 2015; Bittner et al., 2017; Diamantaki et al., 2018; Magee and Grienberger, 2020; Milstein et al., 2021), we first introduced targeted synaptic plasticity exclusively in entorhinal synapses that were active during the target place-field location in the 1D arena (Table 1). This was implemented by increasing AMPA permeabilities of synapses that received inputs from the entorhinal cortex during the target place-field location (Table 1), through synapses distributed across OML and MML dendrites. Across all four target transitions, graded increase in the strength of synapses that were active during the target place-field location expectedly resulted in graded increase in action potential firing within the target location (Fig. 2*B–E*, Δ*P_AMPAR_*). However, such in-field increase was invariably accompanied by spurious increase in firing at other out-of-target-field locations, with spurious firing increasing with increase in synaptic strength. This was particularly pronounced in target transitions that required suppression of preexisting place fields (Fig. 2*D–E*).

For each of these cases, we computed firing rates and spatial selectivity coefficient computed towards assessing the emergence of single place fields (Supplementary Fig. S1*B–C*, Supplementary Fig. S2*B–C*, Supplementary Fig. S3*B–C*, Supplementary Fig. S4*B–C*). These quantifications confirmed an increase in firing rate but a distinct lack of spatial selectivity in the emergent firing profiles, across all four transitions when synaptic changes were the only ones introduced. Such lack of spatial selectivity should be expected because LEC and MEC synapses could have one or two firing fields (Fig. 1*A*), each active within and outside the target arena locations, and a potentiated synapse will increase responses at all these locations where the input is active. Thus, when potentiation altered synapses that were active during the target locations, that also increased firing at other locations by virtue of the entorhinal inputs to these very synapses being active at other off-target locations.

These analyses were performed on the base model, with one set of entorhinal inputs for each of the four transitions and with one specific dendritic localization profile for the afferent synapses. To assess the generality of our findings, we picked one of the four target transitions (stabilization of an existing field) and asked if our conclusions would generalize to 25 different synaptic distributions and to 25 distinct sets of entorhinal inputs (Supplementary Fig. S5). For all these cases, we continued to assess the impact of exclusively introducing graded increases in the strengths of synapses active within the location of an existing place field in the base model. The actual values of firing rates and the spatial selectivity coefficient manifested diversity, depending on the specific distribution or input set employed (Supplementary Fig. S5). However, irrespective of which dendritic localization profile (Supplementary Fig. S5*A*) or which input combinations (Supplementary Fig. S5*B*) we used, we found firing rate to consistently increase and spatial selectivity to consistently decrease with increasing synaptic strength.

We next assessed generalization of these conclusions from the base model with the rest of the valid GC models (Supplementary Fig. S6*A*) with heterogeneous distributions of the underlying ion channels and other parameters (Kumari and Narayanan, 2024). We picked one of the four target transitions (stabilize existing place field) and evaluated the impact of exclusive introduction of synaptic plasticity on firing rate profiles across the arena (Supplementary Fig. S6*B–C*). We used the default dendritic localization profile and default entorhinal input sets (Fig. 2) identically across all 141 models. Even under baseline conditions and default synaptic strength values, the resulting responses were markedly heterogeneous in a manner that was broadly consistent with experimental observations (Zhang et al., 2020). Specifically, whereas some models exhibited strong place-field selectivity, others showed weak responses, and ∼44% remained silent, translating to pronounced model-to-model variability in firing rate as well as in spatial selectivity (Supplementary Fig. S6*B–C*). When we introduced plasticity exclusively in strengths of synapses that were active within the target place-field, firing rate consistently increased, also reducing the fraction of silent cells to ∼9.3% at 10× increase in synaptic strength (Supplementary Fig. S6*B*). However, spatial selectivity consistently reduced with graded increase in synaptic strengths (Supplementary Fig. S6*C*) indicating a lack of sharp tuning to the target field location. These observations collectively confirmed our earlier findings across 25 dendritic localization profiles (Supplementary Fig. S5*A*), 25 entorhinal input sets (Supplementary Fig. S5*B*), and 141 heterogeneous valid models of granule cells (Supplementary Fig. S6*B–C*). In addition, these analyses also underscore the manifestation of pronounced heterogeneity in neural responses, critically dependent on the dendritic localization profile of synapses, the afferent input structure, and the intrinsic properties of the neuron.

Together, these results demonstrate that targeted plasticity confined exclusively to active afferent entorhinal synapses was insufficient to guide selective routing of spatial information towards achieving sharply tuned single-field firing.

### Convergence of plasticity in afferent synaptic and intrinsic mechanisms could achieve target transitions even in the absence of inhibition

If targeted plasticity confined exclusively to active afferent entorhinal synapses was insufficient to achieve single-field firing, what additional plasticity mechanisms could effectuate these target transitions? There are lines of evidence to suggest that suppression of off-field firing is achieved through additional plasticity involving inhibitory synapses that suppress off-target firing (Royer et al., 2012; Grienberger et al., 2017; Robinson et al., 2020; McKenzie et al., 2021; Rolotti et al., 2022; Valero et al., 2022). There are lines of evidence showing that inhibitory conductances are spatially uniform across the arena (Grienberger et al., 2017) and therefore achieve sharply tuned single-field encoding by *globally* suppressing excitation. Specifically, global inhibition not only suppresses out-field excitation, but also limits amplification by other conductances (Hsu et al., 2018) to in-field locations (Grienberger et al., 2017). This is because targeted excitatory plasticity to in-field locations has ensured that with the target locations, the voltage responses are large enough to activate these other conductances despite such global suppression. If *global* suppression of voltage responses is the mechanism that is required to sharpen place-field firing, could this instead be achieved by plasticity to an intrinsic mechanism that does not recruit inhibitory synapses? Could targeted plasticity in afferent synapses be accompanied by intrinsic plasticity mechanisms that *globally suppress* voltage responses towards achieving sharply tuned single-field firing? Could further amplification of in-field responses through additional intrinsic plasticity involving perithreshold ion channels that further *amplify in-field voltage responses*, aid in achieving our target transitions?

Which combinations of *intrinsic conductances* could achieve global suppression of voltage responses while amplifying in-field voltage responses towards achieving our target transitions? There are several possibilities based on the expression profiles of ion-channel conductances in hippocampal granule cells (Supplementary Table S1). However, we were motivated to consider a combination of three specific ion channel conductances that have been shown to express and significantly alter the physiological characteristics of hippocampal granule cells (Crill, 1996; Bender et al., 2003; Ellerkmann et al., 2003; Stegen et al., 2009; Young et al., 2009; Epsztein et al., 2010; Kress et al., 2010; Artinian et al., 2011; Stegen et al., 2012; Mishra and Narayanan, 2021b): the persistent sodium channel (*g_Na_*), the hyperpolarization-activated cyclic nucleotide gated HCN channel (*g_HCN_*), and the inward-rectifying potassium channel (*g_Kir_*).

The primary motivation for picking these three specific ion channels (*g_Na_*, *g_HCN_*, and *g_Kir_*) was the conjunctive plasticity induced in three ion channels by theta-patterned firing (Mishra and Narayanan, 2022), a natural pattern of firing in DG granule cells (Pernia-Andrade and Jonas, 2014; Diamantaki et al., 2016; Zhang et al., 2020). As theta-patterned activity has been strongly associated with synaptic plasticity in the DG (Greenstein et al., 1988; Pavlides et al., 1988; Shors and Dryver, 1994; Beck et al., 2000; Davis et al., 2004; McHugh et al., 2007; Larson and Munkacsy, 2015), we explored the impact of concomitant plasticity in these three ion-channels that underwent long-term increase in conductance values after theta-patterned activity (Mishra and Narayanan, 2022). In addition, the differential subcellular expression profiles of these three ion channels and how they specifically altered neural responses, involving targeted amplification and global suppression, made them attractive choices for exploring conjunctive synaptic and intrinsic plasticity.

Therefore, we hypothesized that these physiologically motivated, conjunctively occurring intrinsic plasticity mechanisms provide the means to locally amplify responses within the target spatial locations and globally suppress firing responses across the arena. In testing this, we probed the possibility that graded plasticity in these four distinct mechanisms, independently and together, could implement the target transitions (Fig. 2*A*) that achieve single-field spatial firing (Fig. 2*B–E*). The four components that underwent plasticity, the ranges of plasticity assessed, the localization of these components within the neuron, the physiological roles of the ion channels, and how each of them is postulated to alter firing in the arena are listed below:

1. **Targeted synaptic plasticity**, implemented as before, by increases in AMPA permeabilities (Δ*P_AMPAR_*: 2–40×) of synapses from the entorhinal cortex distributed across OML and MML dendrites. Targeted synaptic plasticity was implemented exclusively in synapses that were active during the target place-field location in the 1D arena (Table 1).
2. **Axonal intrinsic plasticity**, implemented by increases in sodium conductance (Δ*g_Na_*: 1– 32×) localized to the axon initial segment. Increase in *g_Na_*, a regenerative conductance whose voltage activation is perithreshold in nature, selectively amplifies responses that are large enough to activate these channels while being ineffective on smaller deflections that are subthreshold. These channels strongly regulate granule cell firing rates without altering subthreshold integration or excitability (Mishra and Narayanan, 2021b).
3. **Dendritic intrinsic plasticity**, implemented by increases in HCN conductance (Δ*g_HCN_*: 1–182×) across dendritic compartments. Increase in *g_HCN_*, a resting restorative conductance that acts as a negative feedback loop, globally suppresses responses across the arena. These channels strongly regulate granule cell excitability by altering subthreshold integration and action potential firing (Mishra and Narayanan, 2021b).
4. **Global intrinsic plasticity**, implemented by increases in Kir conductance (Δ*g_Kir_*_1_, Δ*g_Kir_*_2_: 1–2×) globally across all compartments. Increase in *g_Kir_*, a resting conductance that mediates an outward current, globally suppresses responses across the arena. These channels strongly regulate granule cell excitability by altering subthreshold integration and action potential firing (Mishra and Narayanan, 2021b).

While individual plasticity in each of the three ion channels was studied by exclusively altering one channel at a time along with targeted synaptic plasticity, conjunctive plasticity involved simultaneous plasticity in synaptic permeabilities as well as in all three ionic conductances, to different degrees and in disparate combinations (Fig. 2*B–E*; Supplementary Figures S1–S4). Note that for this set of simulations (Fig. 2*B–E*; Supplementary Figures S1– S4), synaptic permeabilities and ion-channel conductances were altered in discrete grade steps. Across all target transitions, independent plasticity of individual conductances coupled to targeted synaptic plasticity yielded expected changes to firing in the arena but did not consistently yield sharply tuned single-field firing (Fig. 2*B–E*; Supplementary Figures S1–S4). Conjunctive increases in the axonal sodium conductance enhanced firing rates but also reduced spatial selectivity owing to an increase in out-field firing as well. Conjunctive increases in dendritic HCN or global Kir conductances globally suppressed excitability, with minimal changes to spatial selectivity (Fig. 2*B–E*; Supplementary Figures S1–S4). Specifically, sodium channel upregulation enhanced overall excitability, while increases in HCN or Kir conductances globally suppressed firing, with marginal changes to spatial selectivity across such independent changes accompanying synaptic plasticity. Strikingly, conjunctive increases in HCN, Kir, and NaP conductances paired with spatially restricted synaptic potentiation, produced robust and selective place-field responses for certain combinations of fold increases (Fig. 2*B–E*). For all target transitions, these observations quantitatively translated to increases in firing rate and spatial selectivity (Supplementary Figures S1–S4). Thus, across all four target transitions, coordinated plasticity across synaptic and intrinsic components enabled selective amplification of target-location responses while suppressing spurious activity elsewhere.

Do these conclusions about conjunctive increases in HCN, Kir, and/or NaP conductances paired with spatially restricted synaptic potentiation generalize to the other valid heterogeneous GC models (Supplementary Fig. S6*A*)? To address this, we evaluated the impact of plasticity in one or all three ion channels occurring together with targeted synaptic plasticity on stabilizing a pre-existing single place field (Supplementary Fig. S6*D–G*). Despite pronounced model-to-model variability transferred from baseline conditions (Supplementary Fig. S6*B–C*) and consistent with prior observations (Fig. 2*B–E*; Supplementary Figures S1– S4), independent sodium channel upregulation along with targeted synaptic plasticity enhanced in-field excitability (Supplementary Fig. S6*D*). In contrast, independent increases in HCN (Supplementary Fig. S6*E*) or Kir (Supplementary Fig. S6*F*) conductances continued to globally suppress firing across all models. However, none of these graded independent increases in ion-channel densities resulted in significant increases in both firing rates and spatial selectivity (Supplementary Fig. S6*D–F*). Strikingly, and again consistent with our observations from the base model (Fig. 2*B–E*; Supplementary Figures S1–S4), conjunctive increases in NaP, HCN, and Kir conductances enhanced firing rates while also maintaining high spatial selectivity with specific combinations of graded synaptic and ion-channel plasticity (Supplementary Fig. S6*G*).

Together, these results demonstrate that conjunctive synaptic and intrinsic plasticity provide a mechanistic basis for the emergence of spatially selective place-cell firing in a heterogeneous granule cell population, even in the absence of inhibitory inputs and without requiring afferent inputs tuned to single spatial locations.

### Exploring the plasticity space to identify combinations that yield sharply tuned single-field place cells

The analyses above provide proof of concept for the ability of conjunctive plasticity in these distinct components (Δ*P_AMPAR_*, Δ*g_Na_*, Δ*g_HCN_*, and Δ*g_Kir_*) in achieving sharply tuned single-place cells after each of the four distinct target transitions (Fig. 2). The analyses were restricted by the discrete grades of plasticity that we had introduced, which were arbitrary and didn’t sample the plasticity space more thoroughly. To address this lacuna, we modified the well-established multi-parametric multi-objective stochastic search (MPMOSS) procedure to stochastically sample the plasticity space, with the objective set at achieving sharply tuned single place fields.

The parameters Δ*P_AMPAR_*, Δ*g_Na_*, Δ*g_HCN_*, Δ*g_Kir_*_1_, and Δ*g_Kir_*_2_were sampled from respective uniform distributions, with Δ*g_Kir_*_1_ and Δ*g_Kir_*_2_representing plasticity in Kir channels expressed in soma/dendrites and AIS/axon, respectively (Table 2). The ranges of each parameter for each target transition were derived from our earlier analyses on the specific ranges that were required to observe successful transitions (Fig. 2*B–E*; Supplementary Figures S1–S4). The randomly chosen plasticity parameters were used to subject the model to that specific combination of synaptic and intrinsic plasticity. Identical entorhinal inputs (as with the pre-plasticity model) were presented to this model with altered permeability/conductances to record the voltage response across the arena. The firing rate within the target (Table 1) place field (*f_PF_*) and the coefficient of spatial selectivity (*C_SS_*) were computed from the recorded voltage responses. The objective associated with the MPMOSS procedure was to ensure the emergence of a sharply tuned place field within a target location (Table 1). Therefore, we declared a specific plasticity combination to be valid if *f_PF_* and *C_SS_* were above specified threshold values (Table 3), to together ensure that the in-field firing was high and the out-field firing was low (Basak and Narayanan, 2018). The ranges of in-field firing rates were set to be consistent with those observed in GC place cells (GoodSmith et al., 2017). For each of the four target transitions (Fig. 2*A*), we independently repeated this entire process for thousands of random samples of the respective plasticity space (Table 2) to identify several plasticity combinations that yielded valid target transitions. We then analyzed all valid plasticity combinations for each target transition, to ask if they were heterogeneous, co-dependent, or clustered. In what follows, we present the results of these independent MPMOSS and analyses procedures, individually for each of the four target transitions (Fig. 2*A*).

### Several non-unique and non-random combinations of synaptic and ion-channel plasticity yielded successful conversion of a silent granule cell to a sharply tuned place cell

We sampled the five-dimensional plasticity space to assess 142,000 unique plasticity combinations to achieve the target transition from a silent cell to a single-field place cell within a pre-specified location (Fig. 3*A*). We found 243 (∼0.17%) plasticity combinations to satisfy the criteria for a sharply tuned place cell, with spatial selectivity well above the set threshold (Fig. 3*B*). Although the plasticity in *g_Na_* required for valid transitions was consistently high (∼60–80×) and that for *g_Kir_*_2_ was consistently low (<4×), there was remarkable diversity in the values of each plasticity parameter that yielded successful transitions (Fig. 3*C*). Importantly, in many models, the fold change in targeted synaptic plasticity was relatively low, implying that strong synaptic plasticity was not always essential in converting silent cells to place cells (Fig. 3*C*).

**Figure 3:**
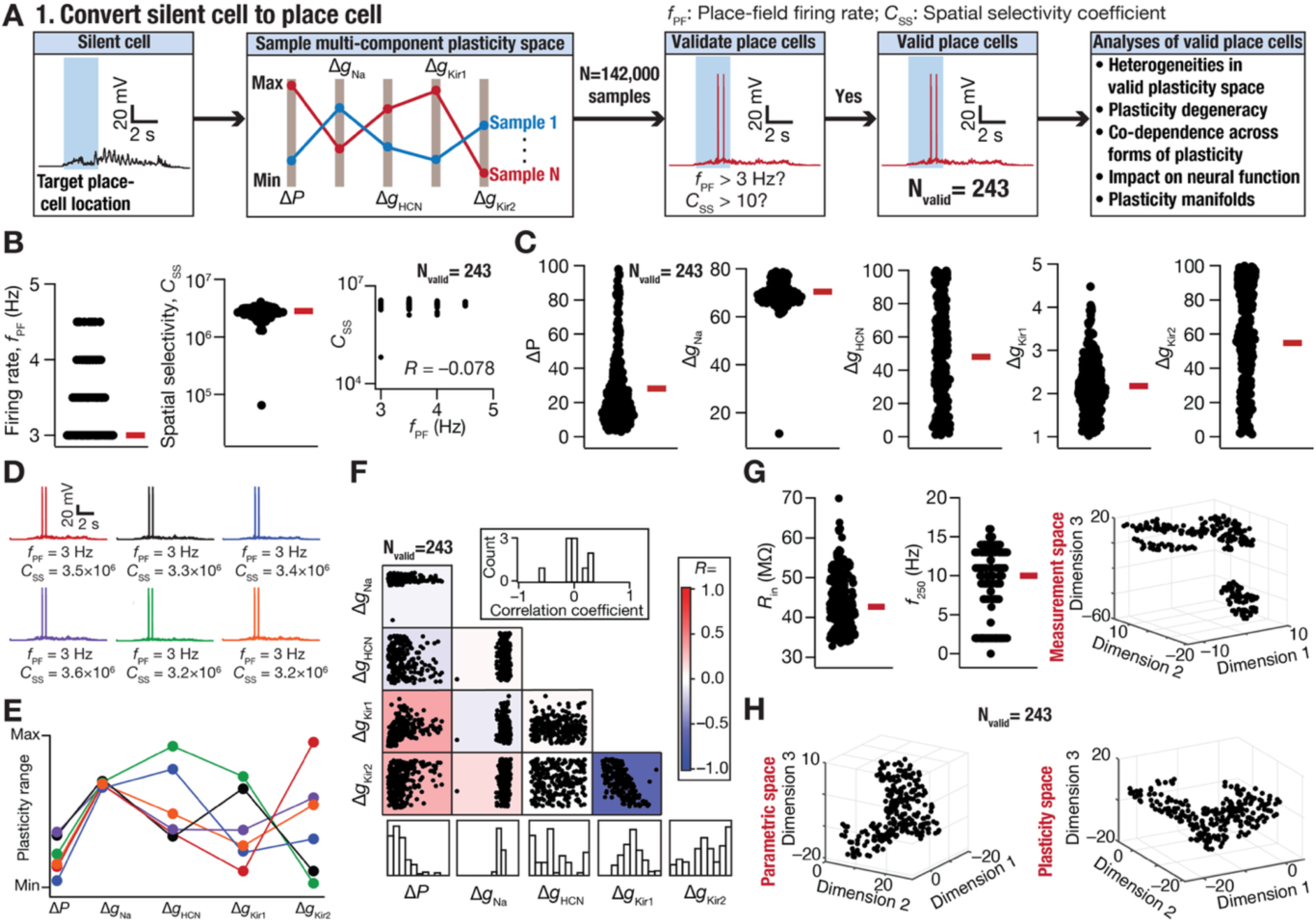
Several non-unique and non-random combinations of conjunctive synaptic and intrinsic plasticity converted a silent DG granule cell to a target specific place cell. (A) Flowchart illustrating the simulation and analyses steps for finding plasticity combinations that converted silent cells to place cells. (B) Beeswarm plots of place-field firing rate (*f*_PF_), spatial selectivity coefficient (*C_ss_*), and the relationship between the two for all the 243 (out of the total 142,000) plasticity combinations that successfully converted a silent cell to a single place-field cell within the targeted region. (C) Beeswarm plots of fold changes in the plasticity parameters from the baseline Δ*P*, Δ*g_Na_*, Δ*g_HCN_*, Δ*g_Kir_*_1_, and Δ*g_Kir_*_2_ for all the 243 valid plasticity combinations. (D) Voltage traces showing place cell response for 6 randomly chosen valid single place-field models with similar *f*_PF_ and *C_ss_* values. (E) Color-coded normalized values of each of the 5 plasticity parameters that yielded the 6 GC place cell models in panel D. (F) Correlation matrix between the five plasticity parameters in all 243 valid models. Heat map shows the correlation values between –1 and +1. Inset shows the histogram of all unique pairwise correlation values across plasticity parameters. Bottom panels show the histograms for each plasticity parameter. (G) *Left*, post-transition input resistance (*R_in_*) and firing rate in response to a 250-pA current injection (*f*_250_) of all 243 valid models. *Right*, 3D plot representing the outcomes of a nonlinear dimensionality reduction technique (*t*–SNE) for the 15-dimensional post-transition measurements space for all 243 models. (H) 3D plots representing the outcomes of *t*–SNE for the 5-dimensional space of actual parametric values after plasticity and 5-dimensional plasticity space involving fold changes, for all 243 models.

We picked six valid models that manifested similar firing profiles across the arena, with very similar *f_PF_* and *C_SS_* as well (Fig. 3*D*). Despite the strong similarity in the functional measurements, the plasticity combinations that yielded the successful transition were diverse (Fig. 3*E*). The observation that only 0.17% of all randomly generated plasticity combinations (Fig. 3*A*) yielded successful transitions demonstrates that the valid plasticity combinations are non-random. In addition and importantly, the diversity in the valid plasticity space (Fig. 3*C– E*) demonstrates non-unique plasticity routes to achieve successful transition from silent cell to place cell. We found that these non-random and non-unique plasticity routes did not exhibit strong pairwise relationships (except for the two Δ*g_Kir_* conductances; *R* = −0.61), exhibiting predominantly weak pairwise correlations among valid plasticity parameters (Fig. 3*F*). The strong negative correlation between the two Δ*g_Kir_* conductances suggests the need to keep the total Δ*g_Kir_* conductance within a limit to avoid enhanced suppression of firing rates.

As intrinsic conductances were subjected to plasticity, we computed the 8 subthreshold and the 9 suprathreshold measurements for all models that underwent successful transitions (Fig. 3*G*, Supplementary Fig. S7). Many of these post-plasticity measurements were expectedly different from their pre-plasticity base model values (Supplementary Tables S2– S3) and were heterogeneously distributed (Fig. 3*G*, Supplementary Fig. S7), reflective of heterogeneity in the valid plasticity space (Fig. 3*C*). To quantify constraints in different spaces, we applied linear and nonlinear dimensionality reduction techniques (PCA, t-SNE, and UMAP) to the 15-dimensional intrinsic measurement space, to the parametric space built with the actual post-plasticity values of 5 parameters (*P_AMPAR_*, *g_Na_*, *g_HCN_*, *g_Kir_*_1_, and *g_Kir_*_2_), and to the plasticity space of fold changes (Δ*P_AMPAR_*, Δ*g_Na_*, Δ*g_HCN_*, Δ*g_Kir_*_1_, and Δ*g_Kir_*_2_) in the five parameters (Fig. 3*G–H*; Supplementary Fig. S8). The measurements and the parametric spaces exhibited low-dimensional subspaces, suggesting that valid place-cell models occupied constrained regions within these spaces. In contrast, the plasticity space showed no such low-dimensional structure (Fig. 3*G–H*; Supplementary Fig. S8), reinforcing the diversity of non-unique routes available to achieve successful transition to a sharply tuned place cell. Thus, despite constraints on post-plasticity intrinsic properties and parameter combinations, the plasticity space remained flexible, highlighting the existence of multiple mechanistic routes for the emergence of spatially selective firing.

Collectively, these results demonstrate that the conversion of silent granule cells into place cells could be achieved through several non-unique and non-random combinations of conjunctive synaptic and intrinsic plasticity, despite the absence of inhibition.

### Several non-unique and non-random combinations of synaptic and ion-channel plasticity yielded successful stabilization of an existing place field in granule cells

For the second transition scenario involving stabilization of an existing place field, we assessed 10,000 unique plasticity combinations and found 325 (3.25%) plasticity combinations to satisfy the criteria for a sharply tuned place cell, with spatial selectivity above the set threshold (Fig. 4*A–B*). We found a strong negative correlation between the place-field firing rate and spatial selectivity coefficient, demonstrating that models with increased place-field firing also manifested a reduction of spatial selectivity. The range of plasticity parameters required for place-field stabilization manifested widespread heterogeneity (Fig. 4*C*). The plasticity ranges for this case were set to smaller ranges compared to the transition from silent cells (Table 2), as increases beyond those ranges resulted in loss of spatial selectivity or place-field firing. Consequently, the range of fold changes here spanned a smaller range (compare with Fig. 3*C*), as the requirement here was a simpler stabilization of an existing field. The transition was nonetheless non-trivial to achieve because merely 3.25% of all randomly generated plasticity combinations (Fig. 4*A*) stabilized the existing place field to yield sharply tuned single-field firing.

**Figure 4:**
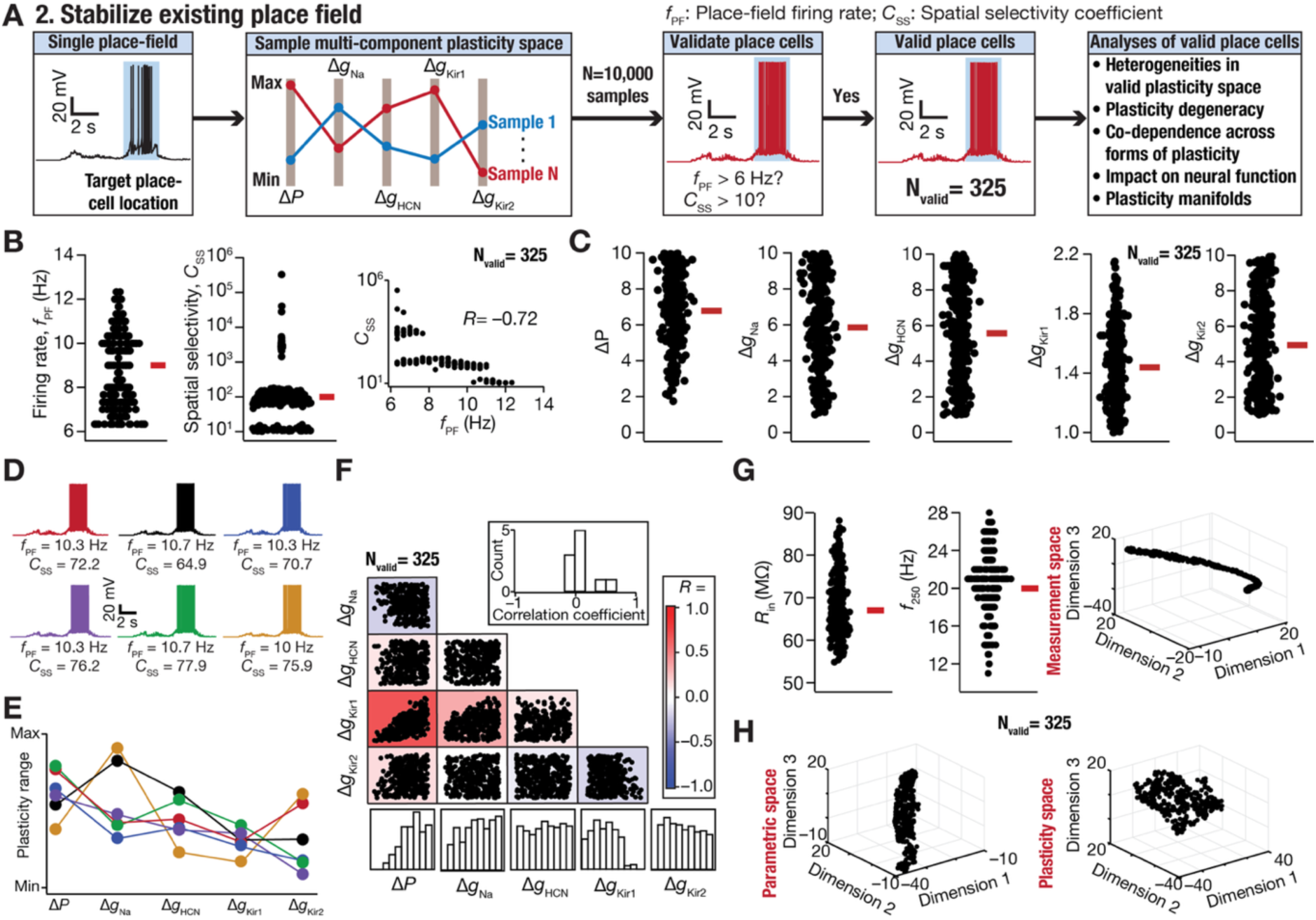
Several non-unique and non-random combinations of conjunctive synaptic and intrinsic plasticity stabilized an existing place field in DG granule cells. (A) Flowchart illustrating the simulation and analyses steps for finding plasticity combinations that stabilized an existing single place field. (B) Beeswarm plots of place-field firing rate (*f*_PF_), spatial selectivity coefficient (*C_ss_*), and the relationship between the two for all the 325 (out of the total 10,000) plasticity combinations that successfully stabilized a single place-field within the targeted region. (C) Beeswarm plots of fold changes in the plasticity parameters from the baseline Δ*P*, Δ*g_Na_*, Δ*g_HCN_*, Δ*g_Kir_*_1_, and Δ*g_Kir_*_2_ for all the 325 valid plasticity combinations. (D) Voltage traces showing place cell response for 6 randomly chosen valid stabilized models with similar *f*_PF_ and *C_ss_* values. (E) Color-coded normalized values of each of the 5 plasticity parameters that yielded the 6 GC place cell models in panel D. (F) Correlation matrix between the five plasticity parameters in all 325 valid models. Heat map shows the correlation values between – 1 and +1. Inset shows the histogram of all unique pairwise correlation values across plasticity parameters. Bottom panels show the histograms for each plasticity parameter. (G) *Left*, post-transition input resistance (*R_in_*) and firing rate in response to a 250-pA current injection (*f*_250_) of all 325 valid models. *Right*, 3D plot representing the outcomes of a nonlinear dimensionality reduction technique (*t*–SNE) for the 15-dimensional post-transition measurements space for all 325 models. (H) 3D plots representing the outcomes of *t*–SNE for the 5-dimensional space of actual parametric values after plasticity and 5-dimensional plasticity space involving fold changes, for all 325 models.

Despite the simplicity of the transition and the reduced range of plasticity search space, six valid models that manifested very similar *f_PF_* and *C_SS_* (Fig. 4*D*) showed heterogeneity in the plasticity combinations that yielded the successful transition (Fig. 4*E*). We found that these non-random (Fig. 4*A*) and non-unique (Fig. 4*C–E*) plasticity routes did not manifest strong pairwise relationships (except for Δ*g_Kir_*_1_*vs*. Δ*P_AMPAR_*; *R* = 0.67), exhibiting predominantly weak pairwise correlations among valid plasticity parameters (Fig. 4*F*). The strong positive correlation between Δ*g_Kir_*_1_and Δ*P_AMPAR_* indicates a coordinated regulation between the two parameters, whereby increases in excitatory synaptic drive were counterbalanced by enhanced potassium conductance to maintain stable single-field firing.

We found post-plasticity intrinsic measurements to be expectedly different from their pre-plasticity base model values (Supplementary Tables S2–S3) and were heterogeneously distributed (Fig. 4*G*, Supplementary Fig. S9), reflective of heterogeneity in the valid plasticity space (Fig. 4*C*). The ranges of these post-plasticity intrinsic measurements were constricted compared to those with the transition of the silent cell (*cf*. Fig. 3*G*, Supplementary Fig. S7), given the restricted plasticity space associated with the stabilization process (Table 2). Dimensionality reduction analyses revealed that the measurements and the parametric spaces exhibited low-dimensional subspaces, while the plasticity space showed no such low-dimensional structure (Fig. 4*G–H*; Supplementary Fig. S10), reinforcing the diversity of routes available for place-field stabilization. These results demonstrate that the stabilization of an existing single-field firing in granule cells could be achieved through several non-unique and non-random combinations of conjunctive synaptic and intrinsic plasticity.

### Several non-unique and non-random combinations of synaptic and ion-channel plasticity yielded successful spatial remapping of an existing place field in granule cells

The third transition requiring spatial remapping of an existing place field is complex compared to the first two transitions because here an existing place field must be suppressed and a sharply tuned place field must emerge at a different target location (Fig. 5). We randomly sampled the plasticity space 5,000 times within the bounds of each of the 5 plasticity parameters (Table 2) and found 139 (∼2.8%) of these samples to satisfy the requirements (Table 3) of the remapping (Fig. 5*A*). The firing rate within the remapped place field and the associated spatial selectivity were above the required threshold (Fig. 5*A–B*), with a negative correlation between *f_PF_* and *C_SS_* across the valid remapped models. The range of plasticity parameters required for place-field remapping spanned a large portion of their respective ranges (Fig. 5*C*, Table 3), despite having merely 2.8% of all randomly generated plasticity combinations remapping the existing place field to an alternate target location.

**Figure 5:**
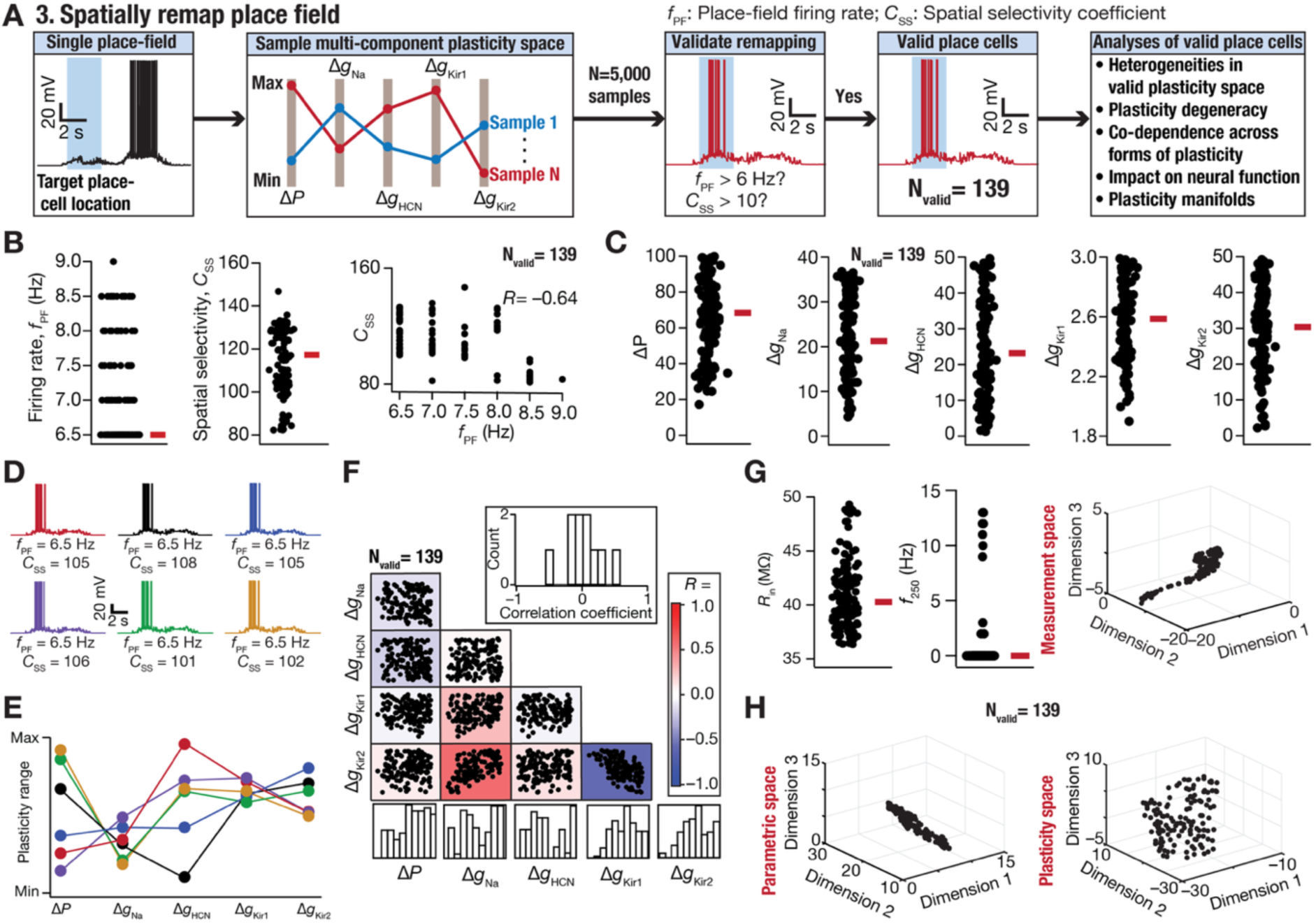
Several non-unique and non-random combinations of conjunctive synaptic and intrinsic plasticity spatially remapped an existing place field in DG granule cells. (A) Flowchart illustrating the simulation and analyses steps for finding plasticity combinations that spatially remapped an existing single place field. (B) Beeswarm plots of place-field firing rate (*f*_PF_), spatial selectivity coefficient (*C_ss_*), and the relationship between the two for all the 139 (out of the total 5,000) plasticity combinations that successfully remapped a single place-field to a targeted region. (C) Beeswarm plots of fold changes in the plasticity parameters from the baseline Δ*P*, Δ*g_Na_*, Δ*g_HCN_*, Δ*g_Kir_*_1_, and Δ*g_Kir_*_2_ for all the 139 valid plasticity combinations. (D) Voltage traces showing place cell response for 6 randomly chosen valid remapped models with similar *f*_PF_ and *C_ss_* values. (E) Color-coded normalized values of each of the 5 plasticity parameters that yielded the 6 remapped models in panel D. (F) Correlation matrix between the five plasticity parameters in all 139 valid models. Heat map shows the correlation values between –1 and +1. Inset shows the histogram of all unique pairwise correlation values across plasticity parameters. Bottom panels show the histograms for each plasticity parameter. (G) *Left*, post-transition input resistance (*R_in_*) and firing rate in response to a 250-pA current injection (*f*_250_) of all 139 valid models. *Right*, 3D plot representing the outcomes of a nonlinear dimensionality reduction technique (*t*–SNE) for the 9-dimensional post-transition subthreshold measurements space for all 139 models. We used only the 9 subthreshold measurements here because only 17 models elicited action potentials, precluding the use of AP measurements for these analyses. (H) 3D plots representing the outcomes of *t*–SNE for the 5-dimensional space of actual parametric values after plasticity and 5-dimensional plasticity space involving fold changes, for all 139 models.

Despite the complexity of the transition requiring a suppression and an emergence, several non-unique plasticity combinations were able to yield successful remapping with very similar *f_PF_* and *C_SS_* (Fig. 5*D–E*). We found that these non-random (Fig. 5*A*) and non-unique (Fig. 5*C–E*) plasticity routes did not require strong pairwise relationships (except for Δ*g_Kir_*_1_*vs*. Δ*g_Kir_*_2_; *R* = −0.56 and Δ*g_Kir_*_2_ *vs*. Δ*g_Na_*; *R* = 0.57 ), exhibiting predominantly weak pairwise correlations among valid plasticity parameters (Fig. 5*F*). The strong positive correlation between Δ*g_Kir_*_2_*vs*. Δ*g_Na_* indicates a coordinated regulation between the regenerative sodium current and the restorative potassium current, whereas the negative relationship between the two Δ*g_Kir_* points to the need for limiting the total *g_Kir_*_1_ in achieving sharply tuned remapping.

We found post-plasticity intrinsic measurements to be expectedly different from their pre-plasticity base model values (Supplementary Tables S2–S3), which were heterogeneously distributed (Fig. 5*G*, Supplementary Fig. S11), reflective of heterogeneity in the valid plasticity space (Fig. 5*C*). Consequent to the requirement to suppress firing within the existing place field, an important difference in post-plasticity intrinsic measurements here compared to the previous two cases (*cf.* Fig. 3*G*, Fig. 4*G*, Supplementary Fig. S7, Supplementary Fig. S9) was the strong suppression of intrinsic excitability (Fig. 5*G*, Supplementary Fig. S11). This suppression manifested as reductions in input resistance, impedance amplitude, temporal summation, and action potential firing rate (Fig. 5*G*, Supplementary Fig. S11). Thus, the range of plasticity here was predominantly defined by the requirement to reduce global excitability towards suppressing an existing firing field. Dimensionality reduction analyses revealed that the measurements space exhibited strongly constricted low-dimensional subspaces, while the parametric and plasticity space showed no such low-dimensional structure (Fig. 5*G–H*; Supplementary Fig. S12), reinforcing the diversity of routes available for place-field remapping.

These results demonstrate that the remapping of an existing place field in granule cells could be achieved through several non-unique and non-random combinations of conjunctive synaptic and intrinsic plasticity. Our observations also highlight the complexity underlying remapping, where the suppression of a pre-existing place field imposed strict requirements involving reduction in intrinsic excitability towards achieving such remapping.

### Several non-unique and non-random combinations of synaptic and ion-channel plasticity yielded successful suppression of spurious second place-field in granule cells

The final form of target transition is another complex transition requiring suppression of one of the two pre-existing place fields while stabilizing the second. Whereas a global suppression would reduce firing rates within both fields, targeted enhancement could still result in non-specific firing beyond the target field that needs to be stabilized (Fig. 6). The percentage of success in finding valid transitions among 50,000 random samples of the plasticity space was 0.45%, resulting in a total of 224 models that suppressed the spurious second field while stabilizing the target field (Fig. 6*A*). There were two subgroups of models, one where the spatial selectivity was high (above ∼300) while in another it was just above the threshold value of 10 (Fig. 6*B*, Table 3). However, there was no correlation between *f_PF_* and *C_SS_* across the valid models (Fig. 6*B*). The range of plasticity parameters required for suppression of the spurious field spanned a large portion of their respective ranges (Fig. 6*C*, Table 3). Whereas the large changes required in sodium channel conductance were comparable to what was required for converting a silent cell to a place cell (*cf*. Fig. 3*C*), changes required in the Kir conductance were also large, with ranges higher than the requirements for remapping (*cf*. Fig. 5*C*).

**Figure 6:**
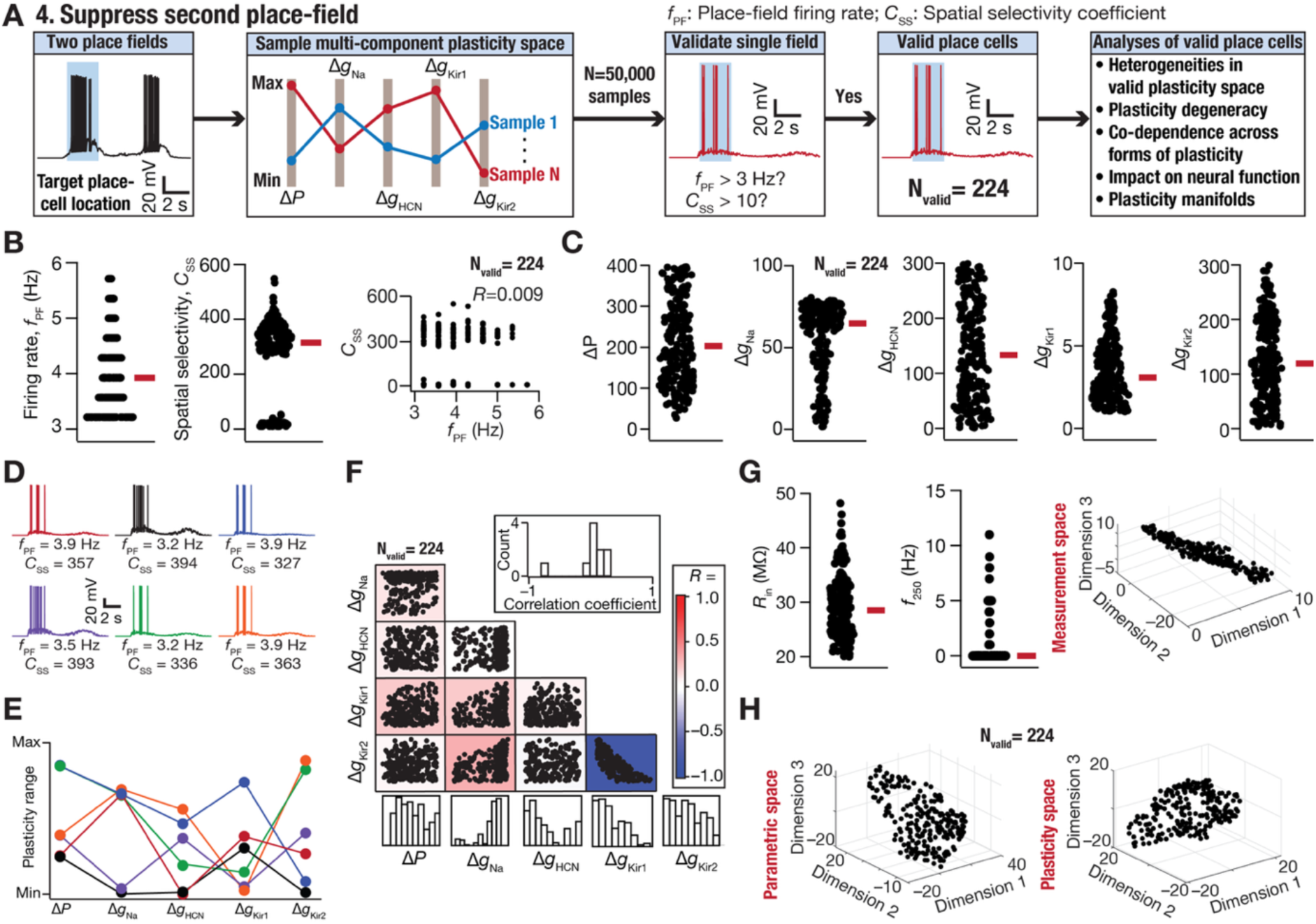
Several non-unique and non-random combinations of conjunctive synaptic and intrinsic plasticity suppressed a spurious second place-field in DG granule cells. (A) Flowchart illustrating the simulation and analyses steps for finding plasticity combinations that suppressed a spurious second place-field. (B) Beeswarm plots of place-field firing rate (*f*_PF_), spatial selectivity coefficient (*C_ss_*), and the relationship between the two for all the 224 (out of the total 50,000) plasticity combinations that successfully suppressed a second place-field beyond the one within the targeted region. (C) Beeswarm plots of fold changes in the plasticity parameters from the baseline Δ*P*, Δ*g_Na_*, Δ*g_HCN_*, Δ*g_Kir_*_1_, and Δ*g_Kir_*_2_ for all the 224 valid plasticity combinations. (D) Voltage traces showing place cell response for 6 randomly chosen valid single place-field models with similar *f*_PF_ and *C_ss_* values. (E) Color-coded normalized values of each of the 5 plasticity parameters that yielded the 6 GC remapped models in panel D. (F) Correlation matrix between the five plasticity parameters in all 224 valid models. Heat map shows the correlation values between –1 and +1. Inset shows the histogram of all unique pairwise correlation values across plasticity parameter. Bottom panels show the histograms for each plasticity parameter. (G) *Left*, post-transition input resistance (*R_in_*) and firing rate in response to a 250-pA current injection (*f*_250_) of all 224 valid models. *Right*, 3D plot representing the outcomes of a nonlinear dimensionality reduction technique (*t*–SNE) for the 9-dimensional post-transition subthreshold measurements space for all 224 models. We used only the 9 subthreshold measurements here because only 38 models elicited action potentials, precluding the use of AP measurements for these analyses. (H) 3D plots representing the outcomes of *t*–SNE for the 5-dimensional space of actual parametric values after plasticity and 5-dimensional plasticity space involving fold changes, for all 224 models.

We found that several non-unique plasticity combinations were able to yield successful suppression of the spurious second field, with very similar *f_PF_* and *C_SS_* (Fig. 6*D–E*), with no requirement for strong pairwise relationships (Fig. 6*F*, except for Δ*g_Kir_*_1_ *vs*. Δ*g_Kir_*_2_; *R* = −0.81). The post-plasticity intrinsic measurements were different from their pre-plasticity base model values (Supplementary Tables S2–S3) and were heterogeneously distributed (Fig. 6*G*, Supplementary Fig. S13). Consequent to the requirement to suppress the spurious second place field, there was strong suppression of intrinsic excitability, manifesting as steep reductions in input resistance, impedance amplitude, temporal summation, and action potential firing rate (Fig. 6*G*, Supplementary Fig. S13). Dimensionality reduction analyses revealed that the measurements space exhibited strongly constricted low-dimensional subspaces, while the parametric and plasticity space showed no such low-dimensional structure (Fig. 6*G–H*; Supplementary Fig. S14). These results demonstrate that the suppression of a second spurious place field in granule cells could be achieved through several non-unique and non-random combinations of conjunctive synaptic and intrinsic plasticity. Our observations also highlight the complexity underlying suppression of a spurious second field, whereby there were strict requirements imposed by the reduction in intrinsic excitability required towards achieving such suppression.

The distinct constraints placed on the specific nature of intrinsic changes required to implement each of the four target transitions emphasize target-dependent variability in the specific forms of plasticity that needs to be accomplished. Specifically, the range of targeted synaptic plasticity and sodium channel plasticity required to create a new place cell or to suppress a spurious field was much higher than that required for stabilization or remapping. The amount of global suppression required through Kir and HCN plasticity was the highest for suppressing the spurious field. The lowest amount of overall plasticity required was to stabilize an existing place field, whereas the highest was to suppress a spurious second field. Consequently, the post-plasticity intrinsic properties showed minimal changes after stabilization, with maximal changes to (reduction) intrinsic excitability occurring after suppression of a second spurious field. The last two target transitions requiring suppression of an existing field achieved this by strong overall reduction of global excitability, although reduction in global excitability played an important role in converting a silent cell to place cell as well.

Together, these results provide a mechanistic framework for understanding how sparse and specific spatial representations can emerge from broadly tuned afferent inputs in the hippocampal circuit. Importantly, our results provide strong lines of evidence that there are diverse routes to achieve sharply tuned single-field firing in hippocampal granule cells, defined by several layers of diversity in inputs, components, targets, forms of plasticity, and strengths of plasticity:

1. The diversity in the specific afferent input patterns arriving onto a granule cell (Fig. 1*A*; Supplementary Fig. S5);
2. The diversity in dendritic localization profiles of these inputs onto a granule cell (Supplementary Fig. S5);
3. The diversity in the molecular composition, including the distribution of ion-channel channels, in the granule cell (Supplementary Fig. S6);
4. The diversity in the naturally emergent (pre-transition) firing responses across the arena (Fig. 1*D*), consequent to diversity in afferent input patterns, dendritic localization, and molecular composition (Fig. 1*A*, Supplementary Figs. S5–S6);
5. The diversity in target-dependent requirements on suppression *vs*. enhancement of firing at specific spatial locations towards achieving sharply tuned single-field firing (Fig. 2), which critically depends on the naturally emergent firing patterns;
6. The diversity in the specific molecular components recruited to implement specific suppression/enhancement required for each transition; and
7. The diversity in the amount of plasticity in each component, defining conjunctive plasticity combinations towards implementing any specific target transition that yields sharply tuned single-field firing (Figs. 3–7).

**Figure 7:**
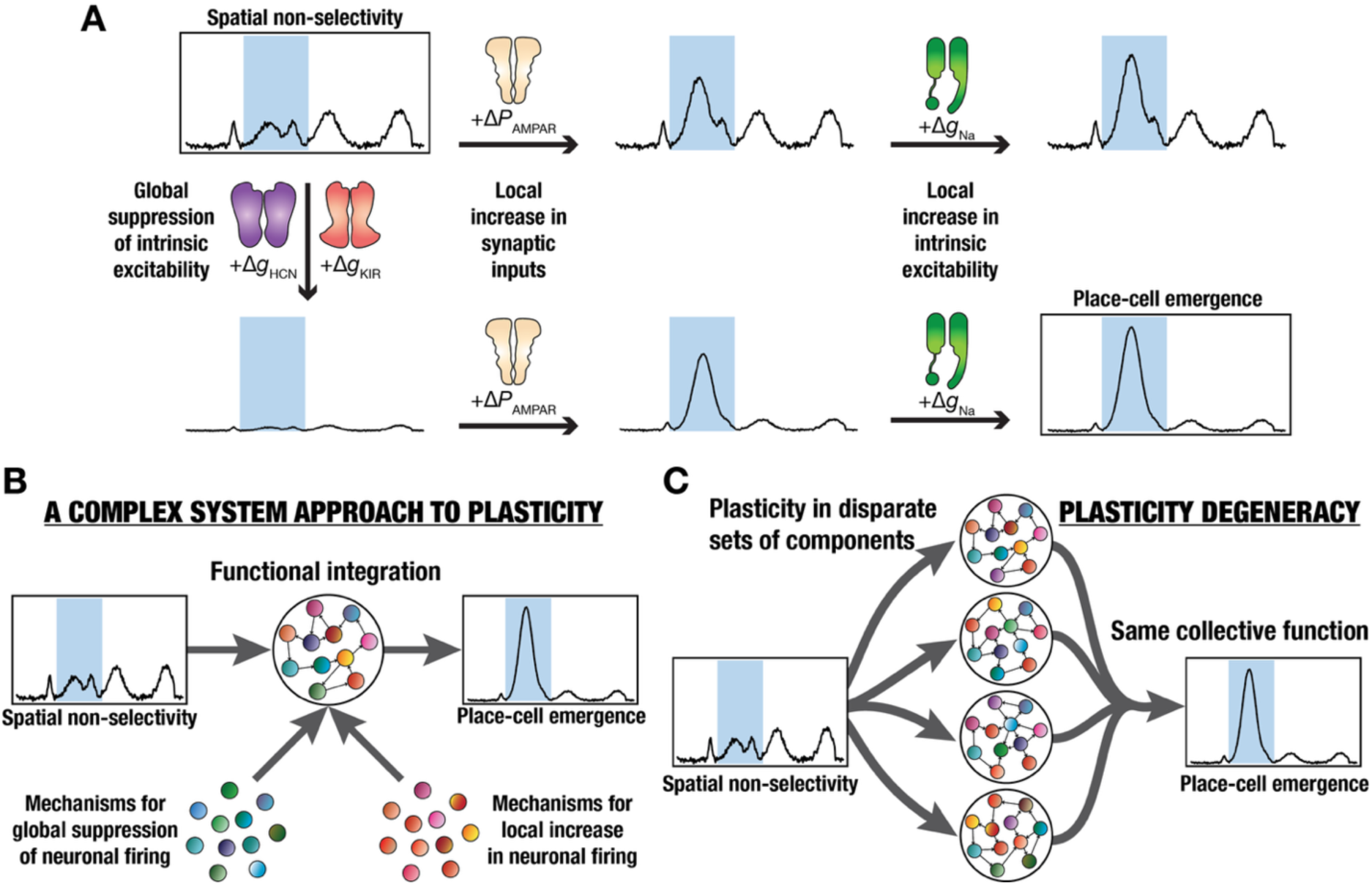
Disparate routes to place-cell emergence in hippocampal granule cells. (A) Schematic illustration of how three distinct plasticity components, mediated by very distinct molecular mechanisms, combined together to yield target transitions that achieved selective spatial routing of spatial information in DG granule cells. The three independent plasticity components are (i) targeted increase in synaptic inputs (mediated by plasticity in AMPAR permeability); (ii) an increase in excitability that selectively amplifies the synaptically amplified voltage response locally because of the voltage-dependent properties (mediated by plasticity in sodium conductances); and (iii) a global reduction in intrinsic excitability that suppresses responses throughout (mediated by plasticity in HCN and Kir conductances). Shown within rectangles are the origin plot where there is spatially non-selective firing (*top left*) and the plot that shows selective place-cell emergence within the target region (*bottom right*). Mere amplification of response in the targeted location, through changes in AMPAR permeability and sodium conductances, does not suppress spurious firing. Mere suppression of intrinsic excitability globally through changes in HCN and Kir conductances does not yield targeted firing. The combination of targeted increase in responses at specific locations and global suppression of intrinsic excitability effectively results in place-cell emergence. (B) There could be disparate mechanisms for global suppression of as well as for local increase in neuronal firing. Such functionally specialized plasticity forms could interact with each other in structured motif-based interactions to yield place-cell emergence from spatial non-selectivity. (C) Disparate combinations of functionally specialized plasticity forms (globally suppressed or locally enhancing neuronal firing) could interact with each other through disparate motif-based interactions to yield place-cell emergence from non-selectivity. The ability of such non-unique and non-random plasticity forms to come together to yield the same collective function represents plasticity degeneracy and underscores the need for a complex system approach to study neural plasticity.

## DISCUSSION

We demonstrate that selective routing of spatial information in dentate gyrus granule cells could emerge through diverse mechanistic routes involving conjunctive plasticity in excitatory synapses and intrinsic ion channels. We show this to be true for emergence of new place fields, for stabilization of existing place fields, for remapping of place fields to alternate location, and for suppression of spurious place fields. Across all target transitions and across a heterogeneous population of models, we provide strong evidence for plasticity degeneracy, wherein disparate combinations of synaptic and intrinsic changes yielded similar routing of spatial information. While distinct transitions warranted different requirements on intrinsic and synaptic properties, the plasticity space remained remarkably flexible, highlighting multiple mechanistic routes to achieve sharply tuned single-field firing in granule cells. These findings highlight that co-dependent plasticity across multiple neuronal components enables robust yet flexible spatial representations, despite substantial heterogeneity in both cellular properties and plasticity mechanisms that were recruited. Together, these findings establish plasticity degeneracy as a key principle underlying flexible formation, robust stabilization, and reliable reconfiguration of spatial representations in granule cells.

### Plasticity degeneracy: A complex system approach to flexible formation and reconfiguration of spatial representations

Our results demonstrate that excitatory synaptic potentiation alone is insufficient to confer sharply tuned spatial specificity to place cells, especially in the DG where the afferent inputs are not from single-field place cells, but the outputs are (Fig. 2; Supplementary Figs. S1–S6; Fig. 7*A*). Thus, the non-specific spatial firing of afferent neurons at multiple spatial locations implies that uniform strengthening of synapses active at any target location results in increased firing but degrades spatial selectivity of action potential firing. These observations highlight a fundamental limitation of targeted excitatory synaptic plasticity in isolation and underscore the need for additional mechanisms to provide suppression of off-field non-specific firing (Fig. 7*A*).

Our choice of specific components to implement local amplification and global suppression was motivated by plasticity in excitatory synapses and specific ion-channel conductances in response to physiologically relevant theta-pattered activity patterns in granule cells (Greenstein et al., 1988; Pavlides et al., 1988; Shors and Dryver, 1994; Beck et al., 2000; Davis et al., 2004; McHugh et al., 2007; Pernia-Andrade and Jonas, 2014; Larson and Munkacsy, 2015; Diamantaki et al., 2016; Zhang et al., 2020; Mishra and Narayanan, 2022).

Driven by these observations from granule cells, we used potentiation in AMPAR to induce targeted changes in specific locations, axonal sodium-channel plasticity for peri-threshold local amplification of potentiated synapses, and plasticity in dendritic HCN and global Kir channels to implement global suppression of firing rates (Fig. 7*A*). Whereas local amplification or global suppression in isolation does not yield well-defined formation or reconfiguration of spatial representations (Fig. 2; Supplementary Figs. S1–S6; Fig. 7*A*), conjunctive plasticity in all components is adequate to effectively achieve sharply-tuned spatial representations.

Our experimental design involved several heterogeneous models of granule cells, several input configurations, several dendritic localization profiles, several target transitions to achieve sharply tuned single fields, and several samples of the 5-dimensional plasticity space offered for each transition. The elaborate nature of our design and the outcomes they yielded, offered an important insight about how neural representations could be stabilized or reconfigured flexibly through several non-unique and non-random combinations of plasticity mechanisms (Figs. 3–6). A straightforward generalization to these findings is that the specific mechanisms that yield global suppression of firing and the ones that implement targeted local amplification need not be from the limited palette of mechanisms that we chose to use in our analyses (Fig. 7*B*). An ideal example for this generalizability is the ability of inhibitory synapses and plasticity therein to implement global suppression of firing in spatially selective neuronal structures (Royer et al., 2012; Grienberger et al., 2017; Robinson et al., 2020; McKenzie et al., 2021; Rolotti et al., 2022; Valero et al., 2022). Global increases in any restorative conductances that are active at rest (*e.g.*, HCN, Kir) or localized increase in active synaptic strengths or peri-threshold regenerative conductances (*e.g.*, voltage-gated sodium or calcium channels, NMDA receptors) could provide supersets of such mechanisms.

If mechanisms that can yield either global suppression or local amplification are considered as *functionally specialized subsystems*, then *functional integration* among these distinct plasticity mechanisms through *synergistic interactions* among them yields collective function (Fig. 7*B*). Importantly, the specific combinations of functionally specialized subsystems are neither completely random nor are completely determined by a unique solution. There are several non-unique and non-random combinations of these functionally specialized subsystems that can yield the collective function of forming and reconfiguring spatial representations. Thus, this plasticity framework presents a non-random interplay between *functional specialization* and *functional integration*, also manifesting plasticity degeneracy where plasticity in disparate structural components yield similar functional outcomes (Fig. 7*C*). Systems with these characteristics are referred to as complex systems (Watts and Strogatz, 1998; Edelman and Gally, 2001; Milo et al., 2002; Kim and Wilhelm, 2008). Thus, based on our results and analyses (Figs. 3–7), we propose that plasticity in neural systems could be analyzed from within a complex systems framework. Within this framework, flexible formation and reconfiguration of neural representations do not arise from a unique plasticity mechanism, but through one of several disparate combinations of functionally specialized components that synergistically interact to yield the required collective function (Fig. 7*C*).

While our analyses have been from the perspective of representations of spatial information, our framework is generalizable to representations of contextual information and their storage within an engram cell perspective (Josselyn and Tonegawa, 2020). Within our framework (Fig. 7), plasticity mechanisms implementing targeted local changes to firing, including excitatory synaptic potentiation and peri-threshold amplification, enhance context-dependent firing. Complementary to this, plasticity mechanisms that implement global suppression, such as inhibition or restorative ion-channel conductances, ensure that neural firing to other contexts is reduced. Together, these complementary forms of plasticity ensure that context-specific firing is implemented by a combination of synaptic and intrinsic (and potentially other) mechanisms. Thus, the approach to achieve stable and flexible neural representations (which can emerge *de novo*, strengthen, weaken, remap, or disappear as animals learn and interact with changing environments) through plasticity degeneracy offers a generalized framework to assess learning and memory through heterogeneous forms of brain plasticity.

Importantly, the complex systems approach to brain plasticity also strongly warns against focusing on any single form of plasticity, as collective function could emerge through the recruitment of one of several diverse combinations of different forms of plasticity. Within the complex system framework, plasticity in any individual component would not yield collective function, and the specific combination chosen would be variable across trials, across animals, and across contexts. Whereas synaptic plasticity could be dominant in one realization of the collective function, plasticity in a specific ion channel would be the dominant form in another realization (*e.g.*, heterogeneities in plasticity space in Figs. 3–6). Thus, singular focus on any individual component or measurements of only one form of plasticity would offer an extremely limited picture of the global plasticity space that yields collective function under different conditions (Kim and Linden, 2007; Titley et al., 2017; Lisman et al., 2018; Josselyn and Tonegawa, 2020; Mishra and Narayanan, 2021a; Mittal and Narayanan, 2024).

### Heterogeneity and degeneracy associated with physiological properties *vs*. plasticity spaces in the hippocampus

Heterogeneities in physiological characteristics and biophysical composition of neurons within the hippocampal formation are electrophysiologically well characterized (Magee and Johnston, 1995; Hoffman et al., 1997; Magee, 1998; Chen and Johnston, 2004; Giocomo et al., 2007; Narayanan and Johnston, 2007, 2008; Narayanan et al., 2010; Cembrowski et al., 2016; Malik et al., 2016; Sun et al., 2017; Cembrowski and Spruston, 2019; Mishra and Narayanan, 2020; Zhang and Jonas, 2020; Mittal and Narayanan, 2022; Kumari and Narayanan, 2026). The ability of these neuronal populations to yield signature electrophysiological characteristics despite widespread heterogeneity in biophysical composition, providing one aspect of cellular-scale degeneracy, has also been very well established (Rathour and Narayanan, 2012, 2014; Beining et al., 2017; Basak and Narayanan, 2018; Migliore et al., 2018; Mittal and Narayanan, 2018; Mishra and Narayanan, 2019; Rathour and Narayanan, 2019; Basak and Narayanan, 2020; Jain and Narayanan, 2020; Mishra and Narayanan, 2020, 2021b; Roy and Narayanan, 2021; Mittal and Narayanan, 2022; Roy and Narayanan, 2022; Kumari and Narayanan, 2024). Furthermore, the ability of disparate molecular components to yield similar short- and long-term stimulus-dependent plasticity profiles in hippocampal neurons has also been demonstrated using detailed biophysical models (Anirudhan and Narayanan, 2015; Mukunda and Narayanan, 2017; Shridhar et al., 2022). Our study takes these lines of research on degeneracy beyond basic physiological properties and plasticity profiles, by demonstrating plasticity degeneracy in yielding similar behaviorally relevant target outcomes through disparate forms of plasticity. We demonstrate plasticity degeneracy to be at the core of yielding *de novo* emergence, stabilization, and reconfiguration of sharply tuned place cells in neurons receiving inputs that are not spatially well-tuned.

While the former demonstrations on degeneracy associated with the characteristic physiological properties allow us to consider a single neuron as a complex system, our analyses here propose a framework that considers neural plasticity as a complex system (Fig. 7). There are other lines of research that have assessed the manifestation of plasticity degeneracy involving different plasticity rules and co-dependent synaptic plasticity in excitatory and inhibitory synapses (Vogels et al., 2011; Agnes and Vogels, 2024; Ramesh et al., 2024; Confavreux et al., 2025a; Confavreux et al., 2025b). Our complex-systems approach to plasticity, which naturally manifests plasticity degeneracy, is a generalized framework involving several forms of plasticity and builds on demonstrations using physiologically constrained models and plasticity spaces involving ion-channel conductances that are known to change together in these cell types. Importantly, our analyses demonstrate that our framework yields the ability to achieve each of *de novo* emergence (Fig. 3), stabilization (Fig. 4), reconfiguration (Fig. 5), and weakening (Fig. 6) of neural representations through non-unique and non-random combinations of plasticity parameters. Together, we argue that the complex systems framework *involving the plasticity space* offers natural explanations of how heterogeneous neural systems implement one of several diverse combinations of plasticity to achieve the precise functional transition required to achieve stable and flexible neural representations.

### Conjunctive plasticity in multiple components is widely prevalent and provides the substrate for disparate plasticity combinations in heterogeneous neural systems

Our observations on how concomitant plasticity in multiple components, each taking neural firing/excitability to opposite directions, together yield target outcomes (Fig. 7) also provide a broader framework to understand experimentally observed scenarios involving such seemingly counter-intuitive combinations. For instance, theta-patterned activity in the dentate gyrus granule cells results in plasticity in synapses and in sodium channels, which enhance excitability in a targeted manner, accompanied by plasticity in Kir and HCN channels, which globally suppress excitability (Greenstein et al., 1988; Pavlides et al., 1988; Shors and Dryver, 1994; Beck et al., 2000; Davis et al., 2004; McHugh et al., 2007; Larson and Munkacsy, 2015; Mishra and Narayanan, 2022). Similarly, theta-patterned activity in CA1 pyramidal neurons yields targeted and localized plasticity in synapses, in *A*-type potassium channels, and in calcium-activated potassium channels, all of which enhance post-synaptic firing, accompanied by a global suppression of excitability through changes in HCN channels (Magee and Johnston, 1997; Frick et al., 2004; Narayanan and Johnston, 2007; Lin et al., 2008; Narayanan and Johnston, 2008). The framework we present here (Fig. 7) cohesively explains the need for such apparently contrasting changes in firing to occur concomitantly towards flexibly achieving well-defined spatial representations. Specifically, mechanisms that suppress global excitability aid in the reduction of off-target firing, while those that provide amplification yield targeted amplification of in-field firing, together ensuring that the specific targets are achieved.

While signaling cascades provide the substrate for *how* such concomitant plasticity in multiple mechanisms could be achieved simultaneously, our framework offers the *functional necessity* for implementing such conjunctive plasticity. These functional constraints also shape the set of plasticity mechanisms that could occur together to define plasticity manifolds, which are structured subspaces within which naturally occurring concomitant plasticity is constrained (Mishra and Narayanan, 2021a; Nagaraj and Narayanan, 2023). For instance, if HCN and Kir channels were to reduce along with an increase in Na and synaptic strength, all plasticity mechanisms would yield increases in excitability, thus resulting in non-specific firing that is detrimental to the targets set in each scenario (Figs. 3–6). Thus, within the framework here, targeted local enhancement of post-synaptic firing must combine with global suppression of excitability to achieve the set goals of selective routing of spatial information. As mentioned above, conjunctive synaptic and intrinsic plasticity observed in hippocampal neurons seems to follow this global principle of local enhancement of firing accompanied by global suppression of excitability through disparate mechanisms. The direction and strength of different forms of plasticity depend on the heterogeneous expression of individual ion channels and the expression/activation of distinct signaling mechanisms that mediate plasticity in each of these components. One of several combinations of such heterogeneous conjunctive plasticity mechanisms together yields specific transitions in a target-dependent manner (Fig. 7). Thus, the overall requirements for conjunctive plasticity in different components with distinct strengths of plasticity, towards implementing degenerate routes towards target transitions, are met by experimental observations where biological plasticity seldom involves single-component plasticity. Future studies could extend our analyses to interactions of plasticity mechanisms among different cell types within the DG and beyond in expanding the systems that are part of such stable and flexible neural representations achieved through plasticity degeneracy (Kim and Linden, 2007; Titley et al., 2017; Lisman et al., 2018; Josselyn and Tonegawa, 2020; Kol and Goshen, 2020; Mishra and Narayanan, 2021a; Mittal and Narayanan, 2024).

In summary, our findings unveil a generalized framework that identifies plasticity degeneracy as a foundational organizing principle underlying the flexible formation and stable reconfiguration of neural representations. Within this framework, flexible memory representations emerge not from specific plasticity rules, but from a repertoire of alternative plasticity routes that are capable of implementing the same computation.

## Author contributions

S.K. and R.N. designed experiments; S.K. performed experiments; S.K. analyzed data; S.K. and R.N. wrote the paper.

## Acknowledgments

The authors thank members of the cellular neurophysiology laboratory for helpful discussions and for comments on a draft of this manuscript.

