## Supplementary figures and tables for "Plasticity degeneracy underlies flexible formation and reconfiguration of spatial representations in hippocampal granule cells"

#### **Supplementary Information**

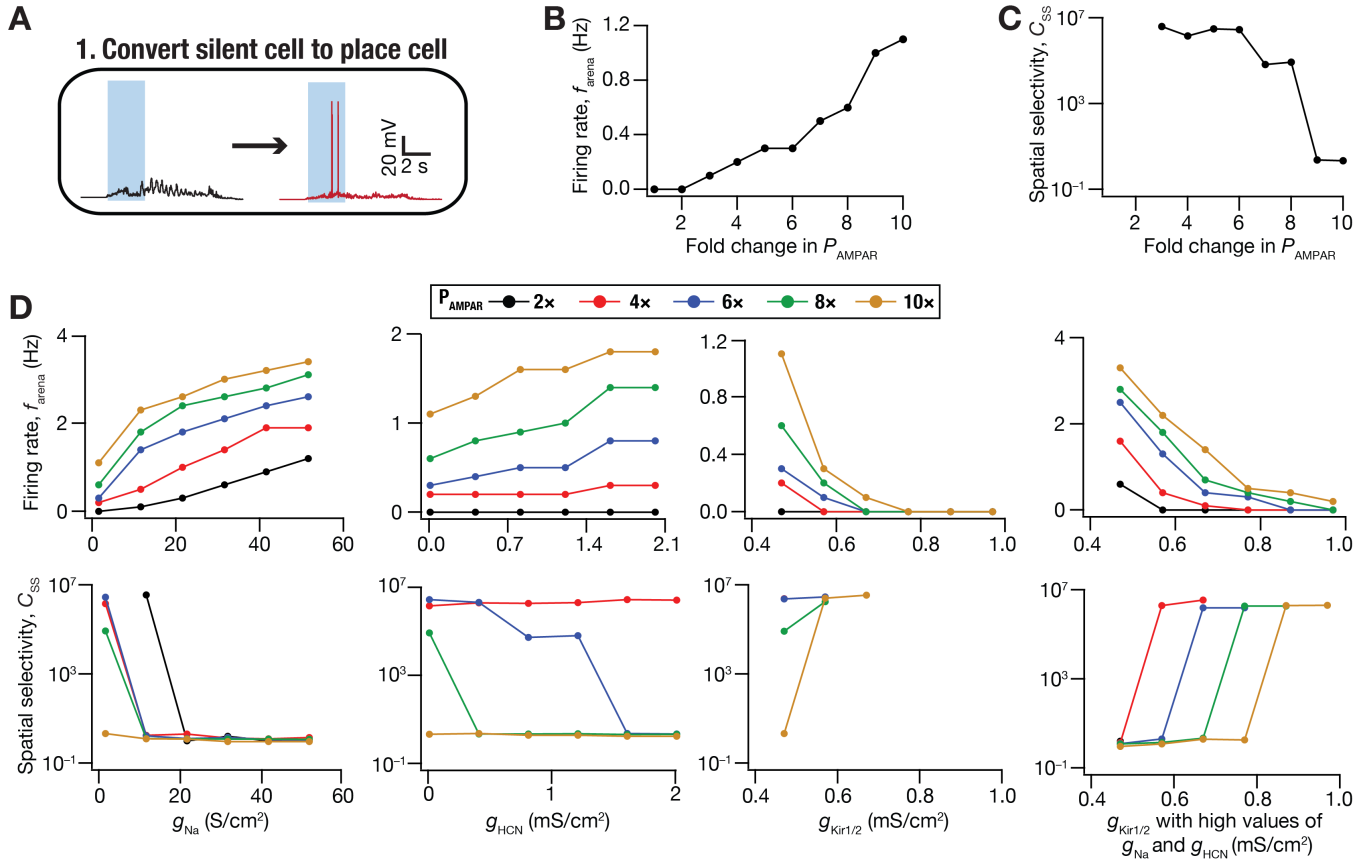

**Supplementary Figure S1: Impact of synaptic and/or intrinsic plasticity in converting a silent DG granule cell to a place cell.** (A) Schematic of the target transition, to convert a silent cell to a place cell, assessed here. The shaded region marks the target location where the place field should emerge. (B–C) Arena firing rate ($f_{\text{arena}}$ in panel B), spatial selectivity coefficient ($C_{\text{SS}}$ in panel C) plotted as functions of fold change in AMPAR permeability ($P_{\text{AMPA}}$). These plots were obtained when firing responses were amplified in the shaded location, exclusively through changes in $P_{\text{AMPA}}$ of synapses carrying spatial information within those locations. (D) Arena firing rate ($f_{\text{arena}}$ in top panels), spatial selectivity coefficient ($C_{\text{SS}}$ in bottom panels) plotted as functions of conductance values of other channels that underwent conjunctive plasticity along with targeted fold changes to $P_{\text{AMPA}}$. The first three column show plots for changes in the conductance values of sodium ($g_{\text{Na}}$), HCN ($g_{\text{HCN}}$), and Kir ($g_{\text{Kir}}$), in that order, accompanying targeted synaptic plasticity. The last column shows plots for conjunctive changes in $P_{\text{AMPA}}$, $g_{\text{Na}}$ (~20× to 30 S/cm<sup>2</sup>), and $g_{\text{HCN}}$ (~10× to 0.1 mS/cm<sup>2</sup>) along with different fold changes in $g_{\text{Kir}}$: $g_{\text{Kir1}}$ in soma and dendrites and $g_{\text{Kir2}}$ in axon and AIS.

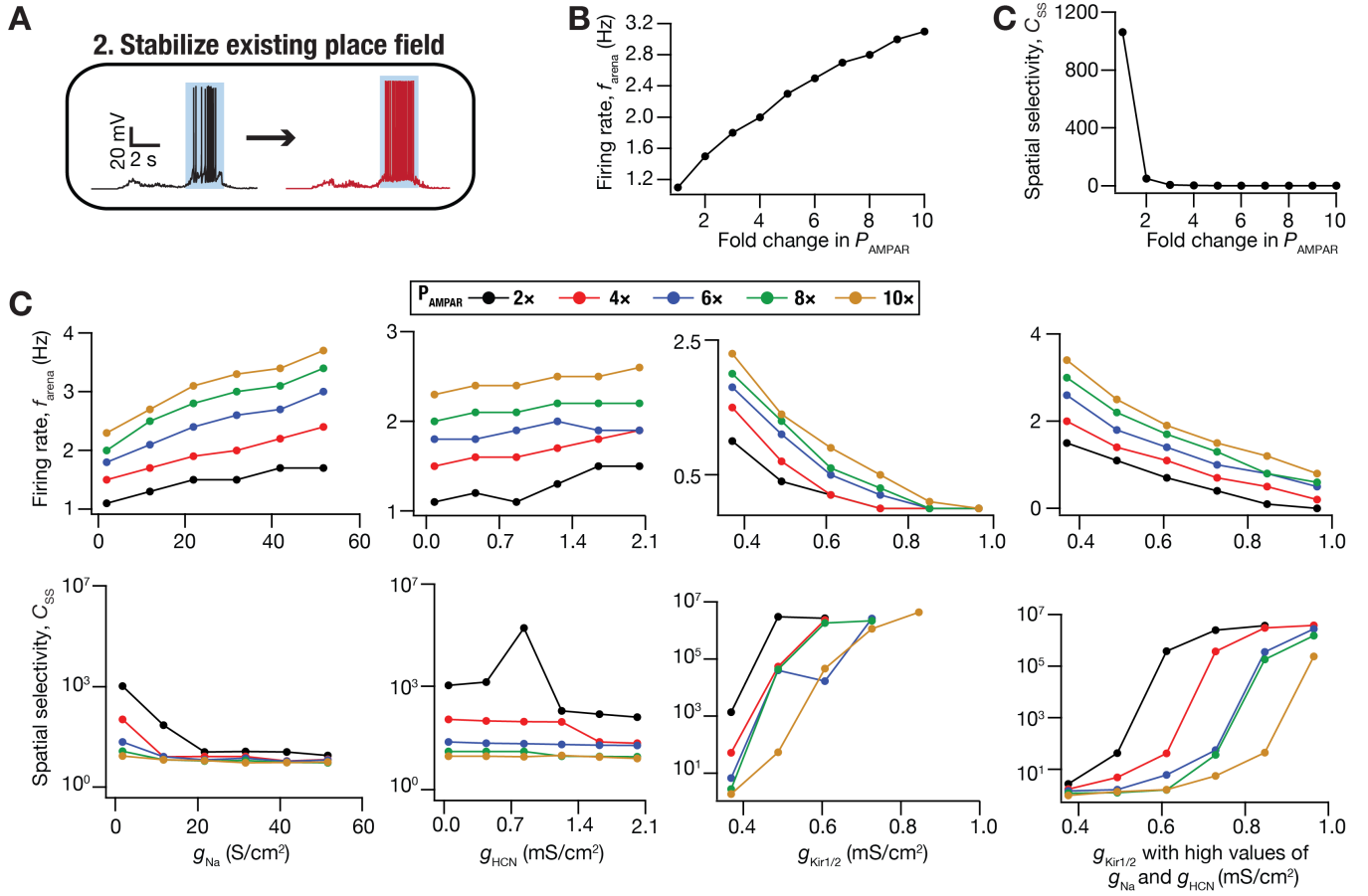

**Supplementary Figure S2: Impact of synaptic and/or intrinsic plasticity in stabilizing an existing DG place cell.** (A) Schematic of the target transition, to stabilize an existing place cell, assessed here. The shaded region marks the target location where the place field should stabilize. (B–C) Arena firing rate ($f_{\text{arena}}$ in panel B), spatial selectivity coefficient ($C_{ss}$ in panel C) plotted as functions of fold change in AMPAR permeability ($P_{\text{AMPAR}}$). These plots were obtained when firing responses were amplified in the shaded location, exclusively through changes in $P_{\text{AMPAR}}$ of synapses carrying spatial information within those locations. (D) Arena firing rate ($f_{\text{arena}}$ in top panels), spatial selectivity coefficient ($C_{ss}$ in bottom panels) plotted as functions of conductance values of other channels that underwent conjunctive plasticity along with targeted fold changes to $P_{\text{AMPAR}}$. The first three column show plots that underwent conjunctive plasticity along with targeted fold changes to $P_{\text{AMPAR}}$. The last column shows plots for conjunctive changes in $P_{\text{AMPAR}}$, $g_{\text{Na}}$ ($\sim 20\times$ to $30$ S/cm$^2$), and $g_{\text{HCN}}$ ($\sim 10\times$ to $0.1$ mS/cm$^2$) along with different fold changes in $g_{\text{Kir}}$. $g_{\text{Kir}}$: $g_{\text{Kir1}}$ in soma and dendrites and $g_{\text{Kir2}}$ in axon and AIS.

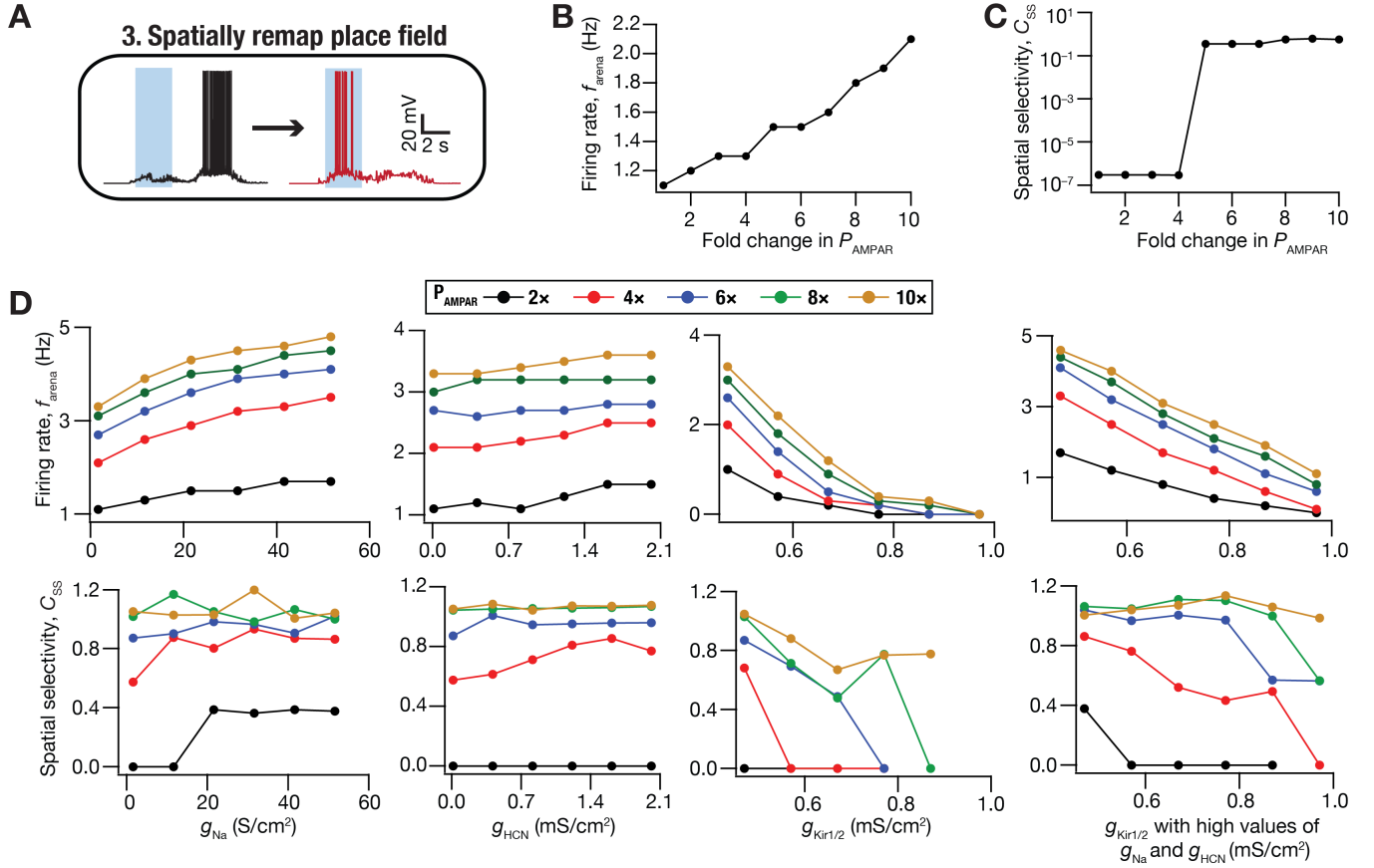

**Supplementary Figure S3: Impact of synaptic and/or intrinsic plasticity in spatially remapping an existing place field in a DG granule cell.** (A) Schematic of the target transition, to spatially remap an existing place field to another location, assessed here. The shaded region marks the target location where the remapped place field should emerge. (B–C) Arena firing rate ($f_{\text{arena}}$ in panel B), spatial selectivity coefficient ($C_{\text{SS}}$ in panel C) plotted as functions of fold change in AMPAR permeability ($P_{\text{AMPAR}}$). These plots were obtained when firing responses were amplified in the shaded location, exclusively through changes in $P_{\text{AMPAR}}$ of synapses carrying spatial information within those locations. (D) Arena firing rate ($f_{\text{arena}}$ in top panels), spatial selectivity coefficient ($C_{\text{SS}}$ in bottom panels) plotted as functions of conductance values of other channels that underwent conjunctive plasticity along with targeted fold changes to $P_{\text{AMPAR}}$. The first three column show plots for changes in the conductance values of sodium ($g_{\text{Na}}$), HCN ($g_{\text{HCN}}$), and Kir ($g_{\text{Kir}}$), in that order, accompanying targeted synaptic plasticity. The last column shows plots for conjunctive changes in $P_{\text{AMPAR}}$, $g_{\text{Na}}$ ($\sim 20\times$ to $30$ S/cm$^2$), and $g_{\text{HCN}}$ ($\sim 10\times$ to $0.1$ mS/cm$^2$) along with different fold changes in $g_{\text{Kir}}$: $g_{\text{Kir1}}$ in soma and dendrites and $g_{\text{Kir2}}$ in axon and AIS.

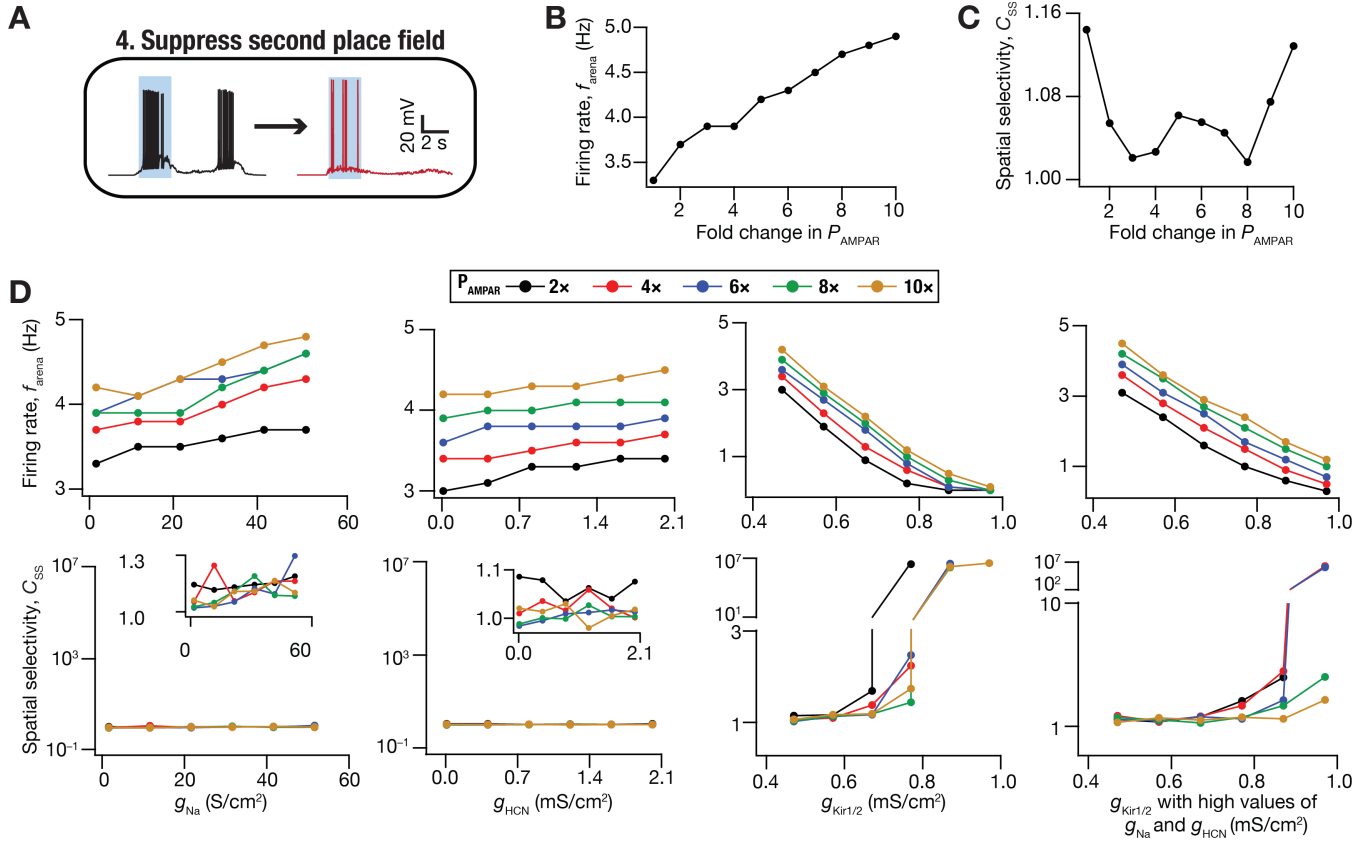

**Supplementary Figure S4: Impact of synaptic and/or intrinsic plasticity in suppressing a spurious second place-field in DG granule cells.** (A) Schematic of the target transition, to suppress a spurious second place-field, assessed here. The shaded region marks the target location where the only place field should manifest. (B–C) Arena firing rate ($f_{\text{arena}}$ in panel B), spatial selectivity coefficient ($C_{\text{SS}}$ in panel C) plotted as functions of fold change in AMPAR permeability ($P_{\text{AMPAR}}$). These plots were obtained when firing responses were amplified in the shaded location, exclusively through changes in $P_{\text{AMPAR}}$ of synapses carrying spatial information within those locations. (D) Arena firing rate ($f_{\text{arena}}$ in top panels), spatial selectivity coefficient ($C_{\text{SS}}$ in bottom panels) plotted as functions of conductance values of other channels that underwent conjunctive plasticity along with targeted fold changes to $P_{\text{AMPAR}}$. The first three column show plots for changes in the conductance values of sodium ($g_{\text{Na}}$), HCN ($g_{\text{HCN}}$), and Kir ($g_{\text{Kir}}$), in that order, accompanying targeted synaptic plasticity. The last column shows plots for conjunctive changes in $P_{\text{AMPAR}}$, $g_{\text{Na}}$ ($\sim 20\times$ to $30$ S/cm$^2$), and $g_{\text{HCN}}$ ($\sim 10\times$ to $0.1$ mS/cm$^2$) along with different fold changes in $g_{\text{Kir}}$. $g_{\text{Kir}}$: $g_{\text{Kir1}}$ in soma and dendrites and $g_{\text{Kir2}}$ in axon and AIS.

**A**

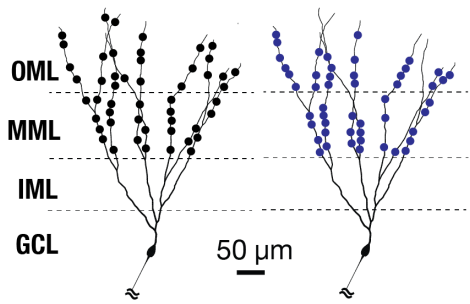

**25 Synaptic distributions**

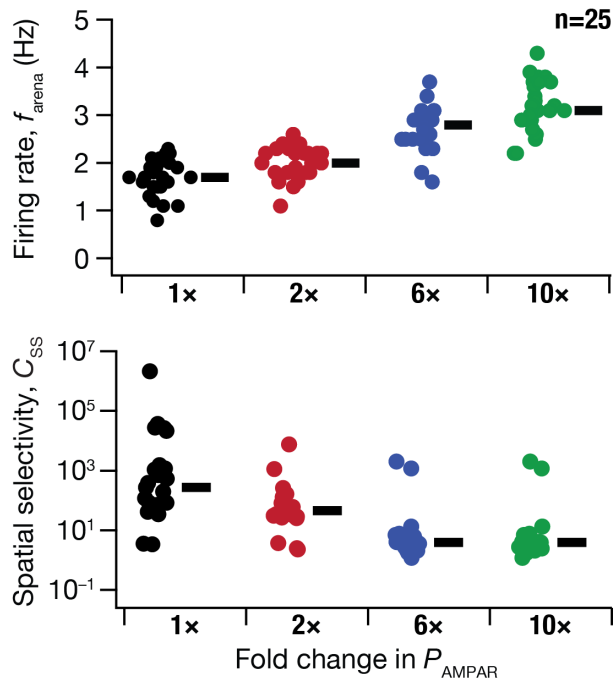

**B**

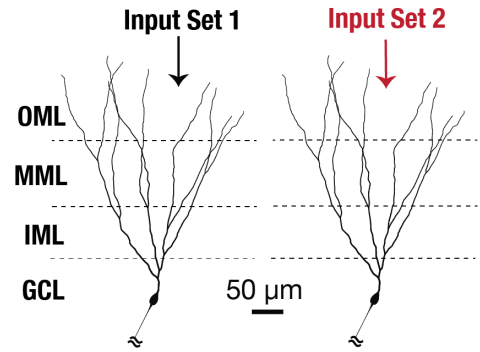

**25 Input sets**

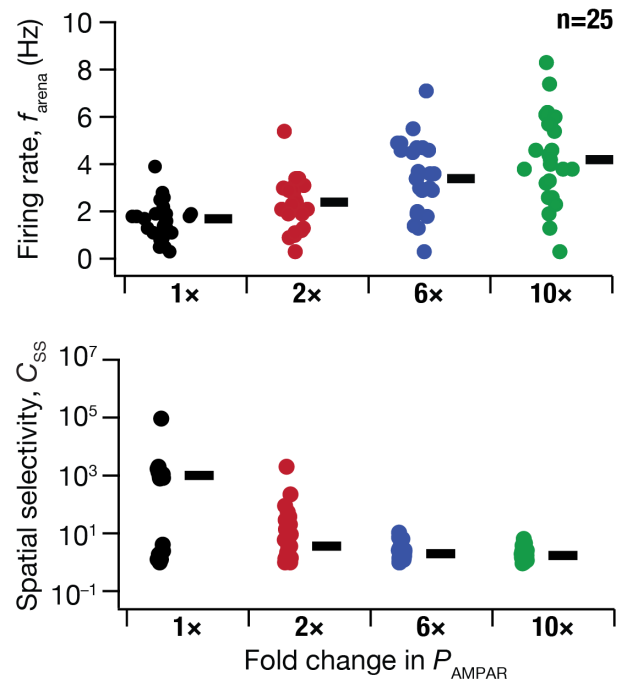

**Supplementary Figure S5: Place field stabilization responses for various synaptic localizations and different entorhinal input sets.** (A) *Top*, schematic for a dendritic tree with different synaptic localization profiles in middle (MML) and outer molecular layer (OML). 25 such localization profiles were generated and received the same set of EC inputs. The target was to stabilize the place field (which was always at the same location owing to the same EC inputs being presented across all synaptic localizations) exclusively through synaptic plasticity. *Middle–Bottom*, arena firing rate (*Middle*) and spatial selectivity coefficient for place fields within the target region (*Bottom*), for each of the 25 localization profiles, plotted as functions of fold changes in synaptic permeability in the targeted region (B) *Top*, schematic for two dendritic trees with different sets of EC inputs impinging on the middle (MML) and outer molecular layer (OML) with the same synaptic localization profiles. 25 such distinct input profiles were generated and elicited place cells at different locations each time. The target was to stabilize the place field exclusively through synaptic plasticity. *Middle–Bottom*, arena firing rate (*Middle*) and spatial selectivity coefficient for place fields within the target region (*Bottom*), for each of the 25 input sets, plotted as functions of fold changes in synaptic permeability in the targeted region.

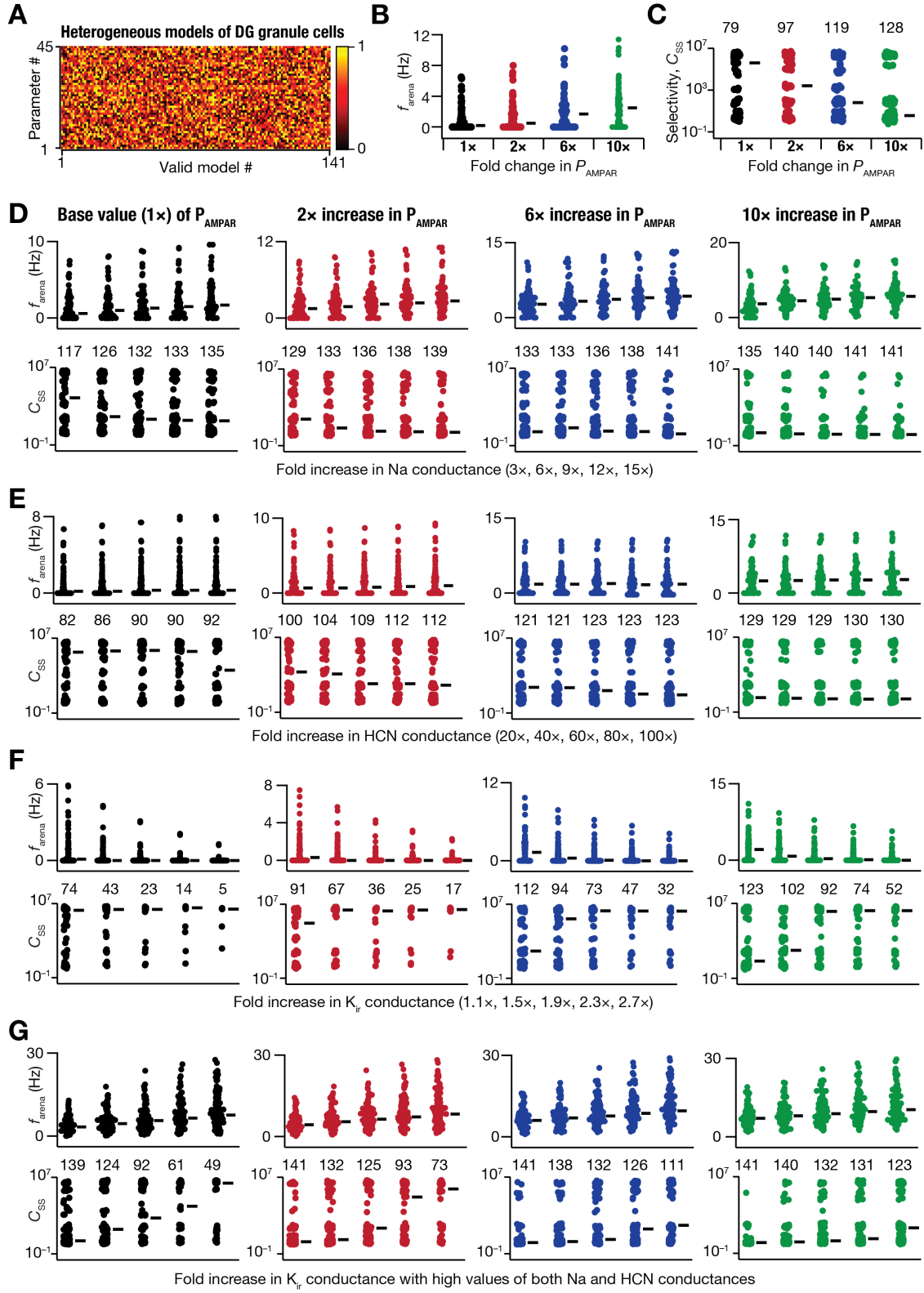

**Supplementary Figure S6: Quantification of place cell response from a heterogeneous population of dentate gyrus granule cell models stabilizing an existing place field through conjunctive synaptic and**

**intrinsic plasticity.** (A) Heat map showing the heterogeneous 45-dimensional parametric distribution of 141 dentate gyrus granule cell models (from Kumari and Narayanan, *J Neurophysiology*, 2024). (B–C) Plots of firing rate for the arena (B) and spatial selectivity coefficient (C) were computed for various fold increases in AMPAR permeability of synapses carrying targeted spatial information, across all 141 models from panel A. For these plots, firing responses were amplified in the targeted location, exclusively through changes in $P_{AMPAR}$ of synapses carrying spatial information within locations where the existing place field was to be stabilized (see Fig. 2C). (D–G) Arena firing rate ($f_{arena}$ in top panels), spatial selectivity coefficient ($C_{ss}$ in bottom panels) plotted as functions of conductance values of other channels that underwent conjunctive plasticity along with targeted fold changes to $P_{AMPAR}$ at different levels (Columns 1–4). These analyses were performed for all 141 models from panel A, for different changes in the conductance values of sodium, $g_{Na}$ (D), HCN $g_{HCN}$ (E), and Kir, $g_{Kir}$ (F) channels, individually accompanying targeted synaptic plasticity. These analyses were also repeated for all models with conjunctive changes in $P_{AMPAR}$, $g_{Na}$ ($\sim 20\times$ to $30$ S/cm<sup>2</sup>), and $g_{HCN}$ ($\sim 10\times$ to $0.1$ mS/cm<sup>2</sup>) along with different fold changes in $g_{Kir}$ (same fold changes as in panel F). $g_{Kir}$: $g_{Kir1}$ in soma and dendrites and $g_{Kir2}$ in axon and AIS. All firing rates are plotted for all 141 models, but the spatial selectivity values are plotted only for those models which elicited non-zero action potentials. Across panels C–G, the numbers above spatial selectivity distributions represent the number of models that elicited non-zero action potentials for that particular case. The thick lines represent the median of the respective distribution.

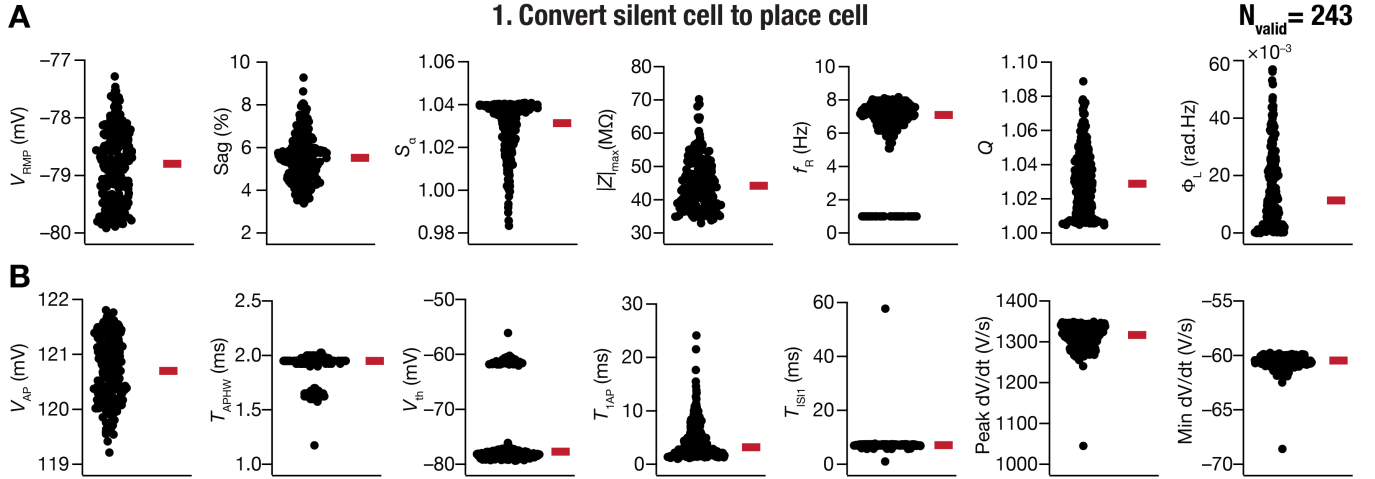

**Supplementary Figure S7: Heterogeneous distribution of electrophysiological measurements of valid DG GC models that successfully converted silent cells to place cells.** (A–B) Beeswarm plots for intrinsic subthreshold (A) and supratherapeutic (B) of the 243 models of DG GC that successfully converted silent cells to place cells through conjunctive intrinsic and synaptic plasticity (Fig. 3). The red rectangle adjacent to each plot depicts the respective median values. $f_{50}$ for all the models was zero and is not plotted here.

### 1. Convert silent cell to place cell

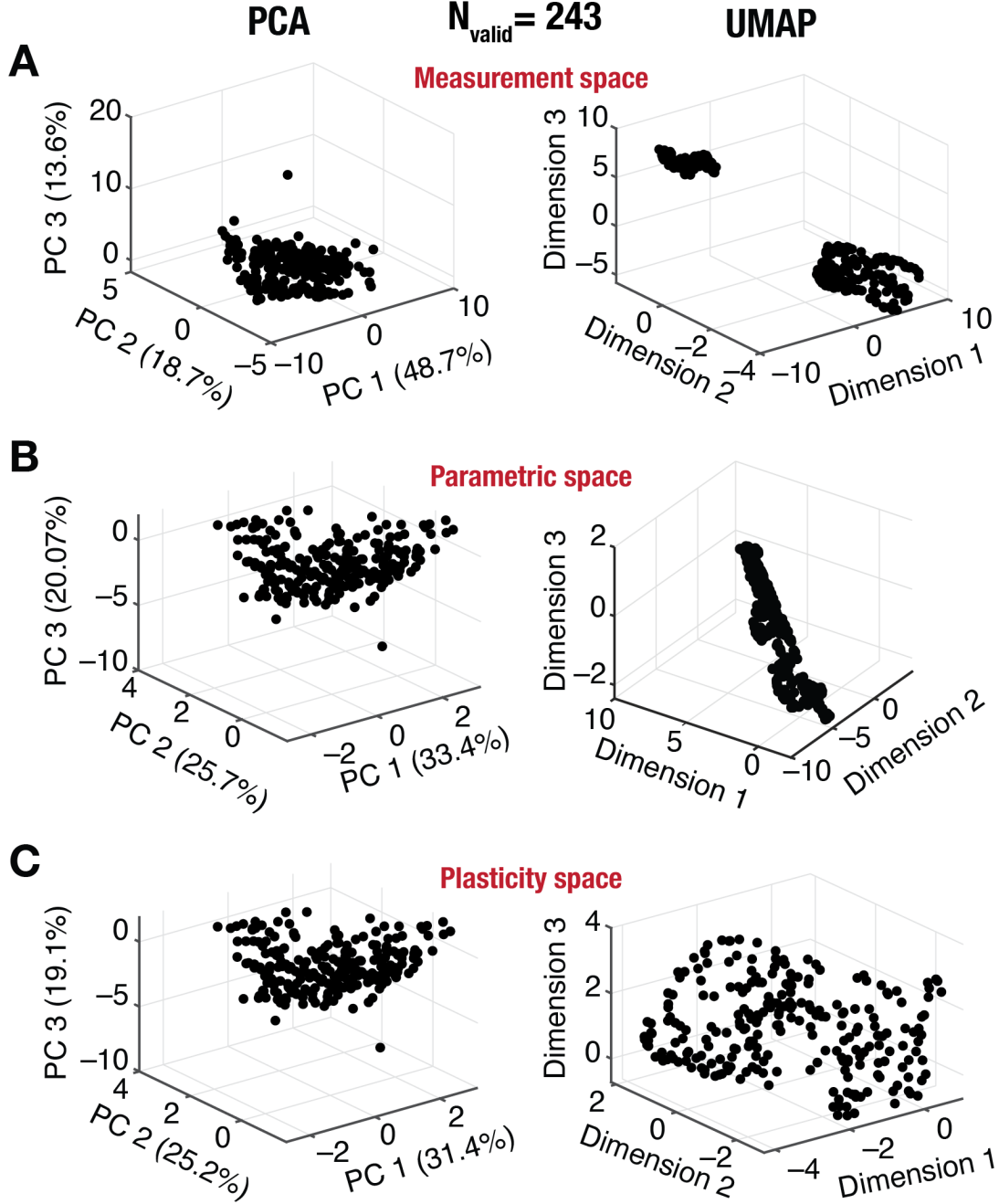

**Supplementary Figure S8: Plasticity degeneracy in converting a silent DG granule cell to a target specific place cell through conjunctive synaptic and intrinsic plasticity.** (A–C) 3D plots representing the outcomes of a linear (PCA, Left column) and a non-linear (UMAP, Right column) dimensionality reduction technique for 15-dimensional measurements space (A), 5-dimensional space of actual parametric values after plasticity (B), and 5-dimensional plasticity space involving fold changes (C), plotted in that order for all 243 valid models (see Figure 3G–H). The variance explained by PC1, PC2 and PC3 are mentioned in the respective axes of all PCA plots. The effective dimensionality values (computed as $(\sum \lambda_i)^2 / \sum \lambda_i^2$, for all eigen values $\lambda_i$) computed from PCA was 3.38, 3.95, and 3.95 for the measurements, parametric, and plasticity spaces, respectively.

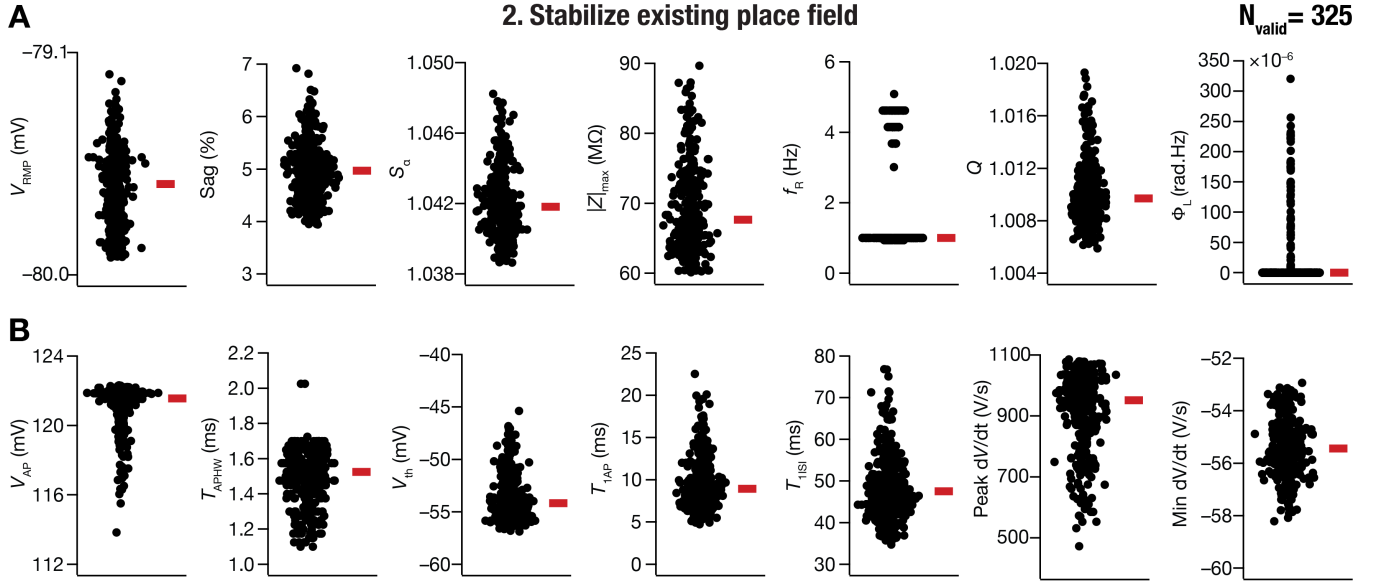

**Supplementary Figure S9: Heterogeneous distribution of electrophysiological measurements of valid DG GC models that successfully stabilized an existing place field.** (A–B) Beeswarm plots for intrinsic subthreshold (A) and suprathreshold (B) measurements of the 325 models of DG GC that successfully stabilized an existing place field through conjunctive intrinsic and synaptic plasticity (Fig. 4). The red rectangle adjacent to each plot depicts the respective median values. $f_{50}$ for all the models was zero and is not plotted here.

#### 2. Stabilize existing place field

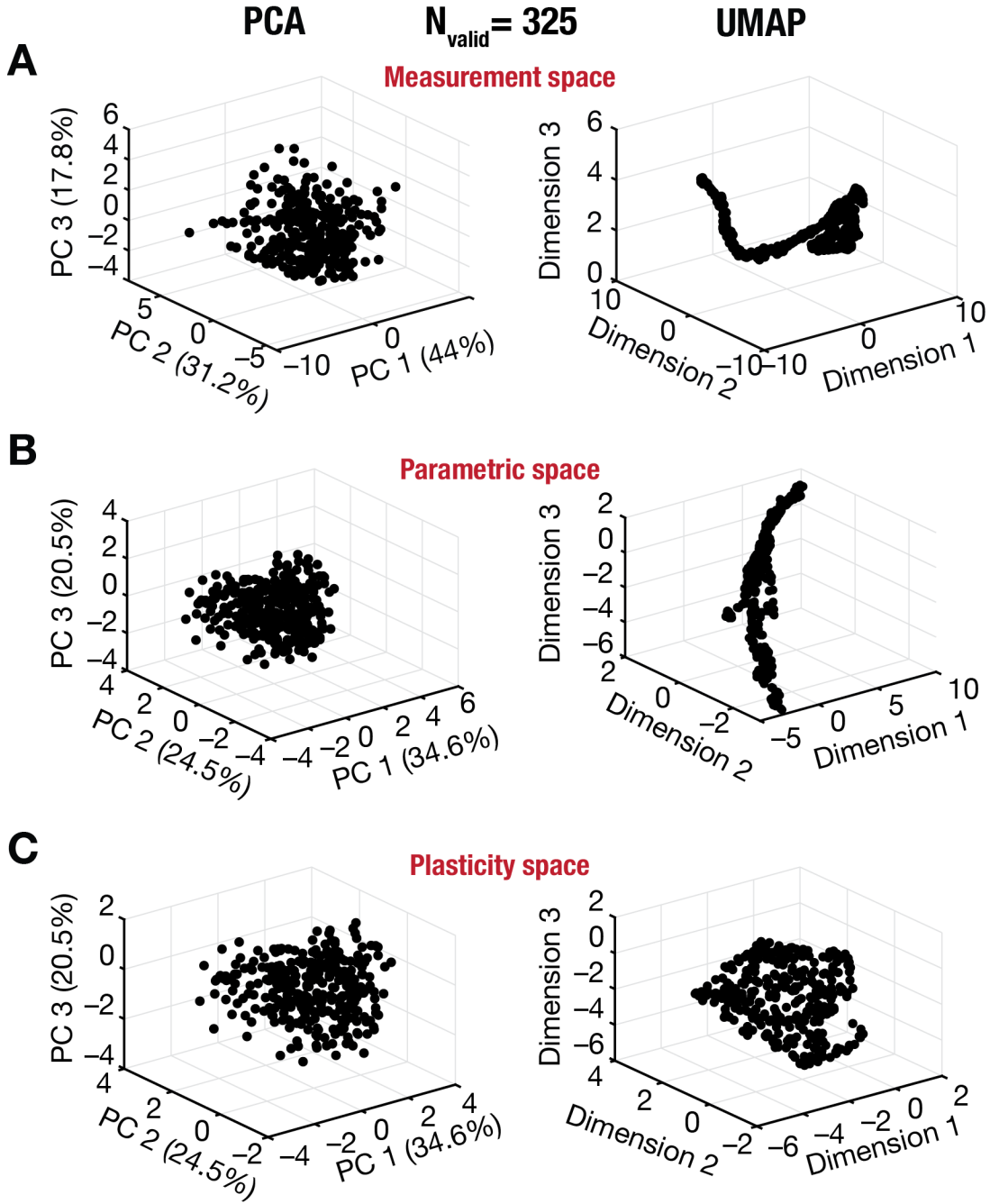

**Supplementary Figure S10: Plasticity degeneracy in stabilizing an existing place field in DG granule cells through conjunctive synaptic and intrinsic plasticity.** (A–C) 3D plots representing the outcomes of a linear (PCA, Left column) and a non-linear (UMAP, Right column) dimensionality reduction technique for 15-dimensional measurements space (A), 5-dimensional space of actual parametric values after plasticity (B), and 5-dimensional plasticity space involving fold changes (C), plotted in that order for all 325 valid models (see Figure 4G–H). The variance explained by PC1, PC2 and PC3 are mentioned in the respective axes of all PCA plots. The effective dimensionality values (computed as $(\sum \lambda_i)^2 / \sum \lambda_i^2$, for all eigen values $\lambda_i$) computed from PCA was 3.2, 3.9, and 3.9 for the measurements, parametric, and plasticity spaces, respectively.

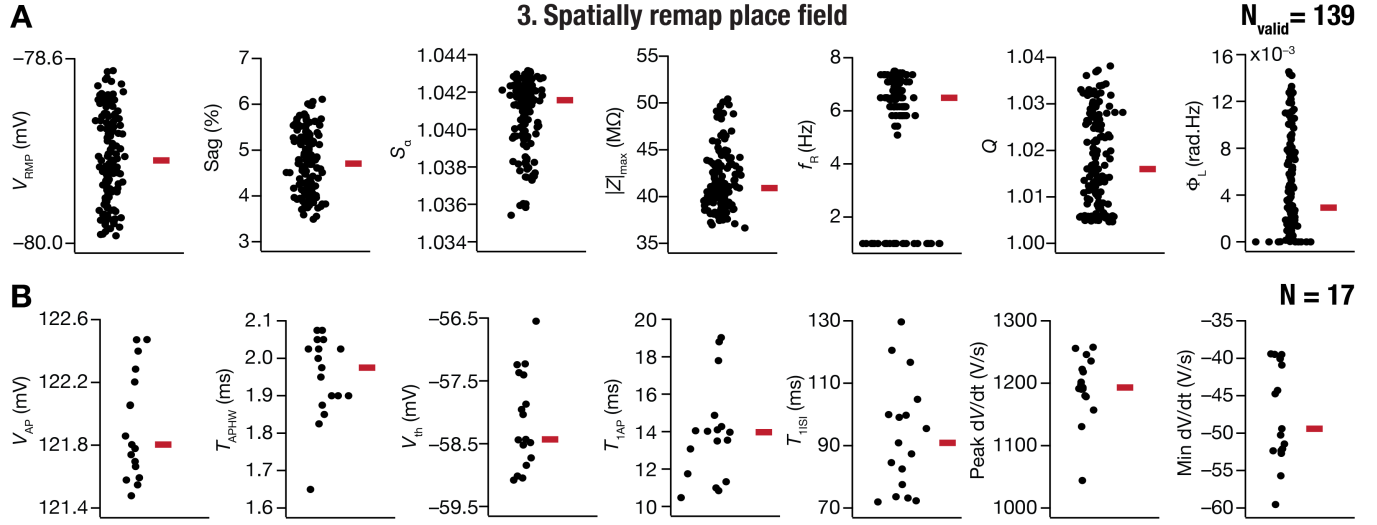

**Supplementary Figure S11: Heterogeneous distribution of electrophysiological measurements of valid DG GC models that successfully remapped an existing place field.** (A–B) Beeswarm plots for intrinsic subthreshold (A) and suprathreshold (B) measurements of the 139 models of DG GC that successfully remapped an existing place field through conjunctive intrinsic and synaptic plasticity (Fig. 5). The red rectangle adjacent to each plot depicts the respective median values. $f_{50}$ for all the models was zero and is not plotted here. The $n$ values in panel B correspond to number of models where action potential firing was observed for 250-pA current injection.

##### 3. Spatially remap place field

PCA

$N_{\text{valid}} = 139$

UMAP

**A**

Measurement space

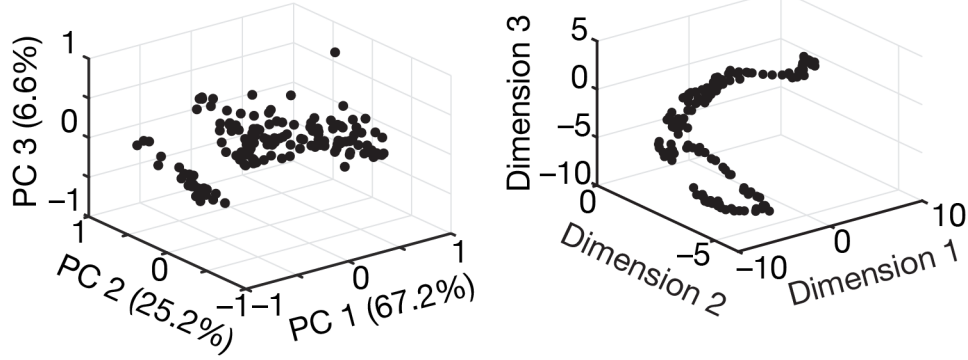

**B**

Parametric space

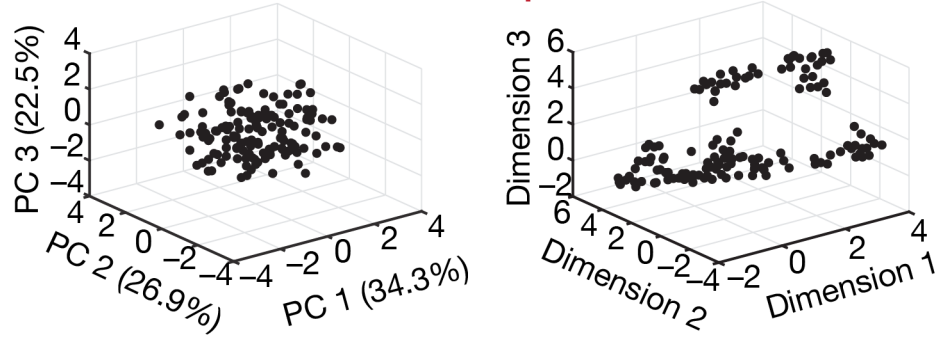

**C**

Plasticity space

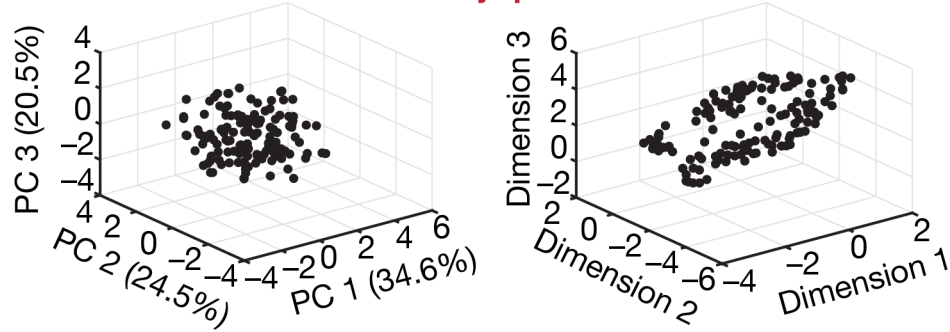

**Supplementary Figure S12: Plasticity degeneracy in spatially remapping an existing place field in DG granule cells through conjunctive synaptic and intrinsic plasticity.** (A–C) 3D plots representing the outcomes of a linear (PCA, Left column) and a non-linear (UMAP, Right column) dimensionality reduction technique for 9-dimensional subthreshold measurements space (A), 5-dimensional space of actual parametric values after plasticity (B), and 5-dimensional plasticity space involving fold changes (C), plotted in that order for all 139 valid models (see Figure 5G–H). The variance explained by PC1, PC2 and PC3 are mentioned in the respective axes of all PCA plots. The effective dimensionality values (computed as $(\sum \lambda_i)^2 / \sum \lambda_i^2$, for all eigen values $\lambda_i$) computed from PCA was 2.5, 3.78, and 3.78 for the measurements, parametric, and plasticity spaces, respectively.

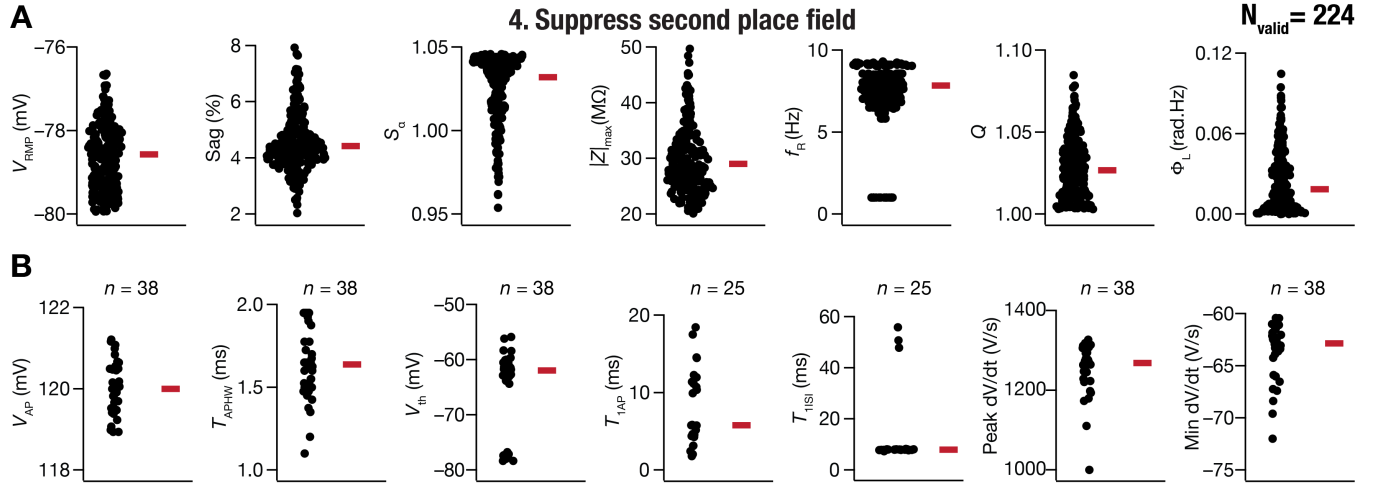

**Supplementary Figure S13: Heterogeneous distribution of electrophysiological measurements of valid DG GC models that successfully suppressed a spurious second place field.** (A–B) Beeswarm plots for intrinsic subthreshold (A) and suprathreshold (B) of the 224 models of DG GC that successfully suppressed a spurious second place field through conjunctive intrinsic and synaptic plasticity (Fig. 6). The red rectangle adjacent to each plot depicts the respective median values. $f_{50}$ for all the models was zero and is not plotted here. The $n$ values in panel B correspond to number of models where action potential firing was observed for 250-pA current injection.

#### 4. Suppress second place field

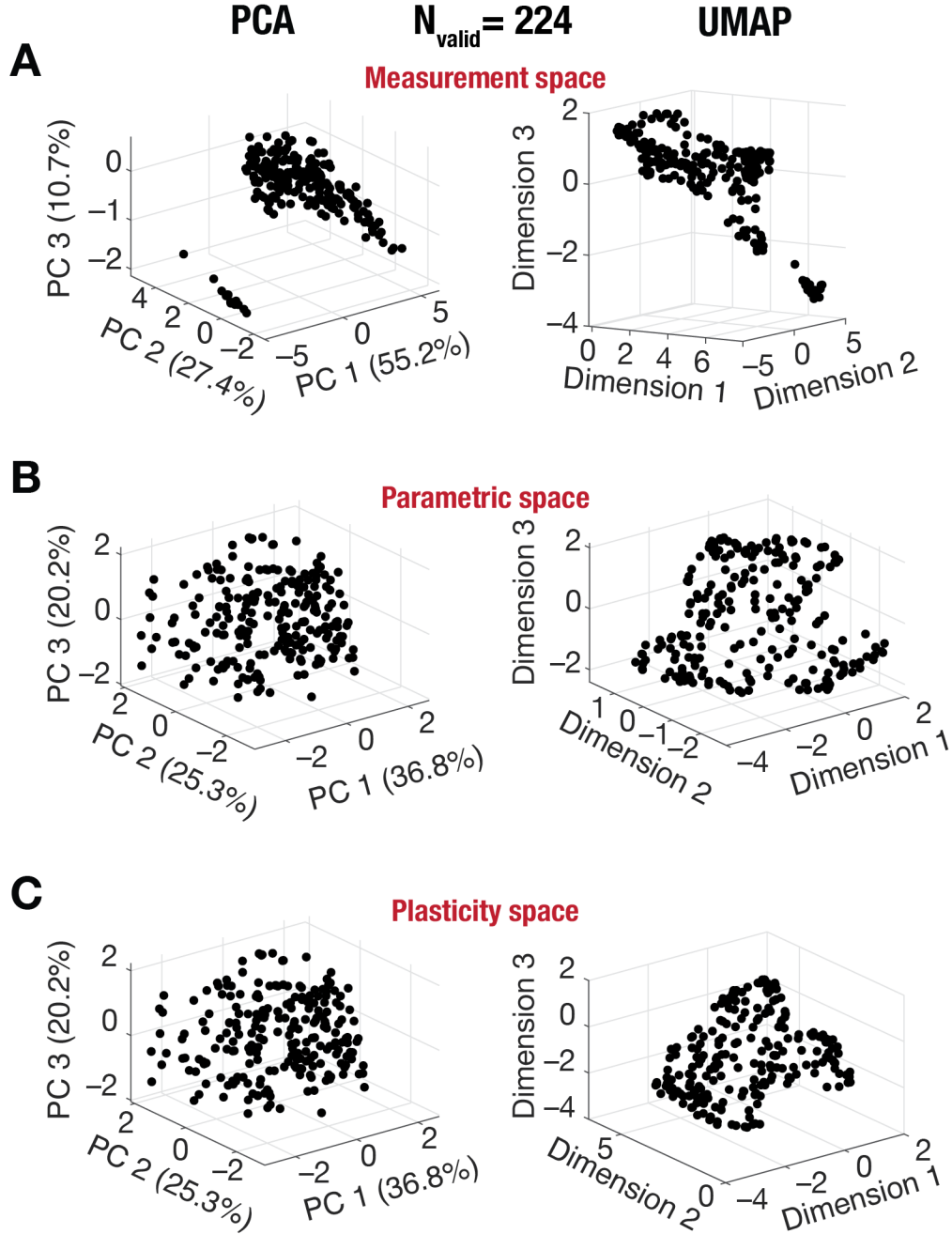

**Supplementary Figure S14: Plasticity degeneracy in suppressing a spurious second place field in DG granule cells through conjunctive synaptic and intrinsic plasticity.** (A–C) 3D plots representing the outcomes of a linear (PCA, Left column) and a non-linear (UMAP, Right column) dimensionality reduction technique for 15-dimensional measurements space (A), 5-dimensional space of actual parametric values after plasticity (B), and 5-dimensional plasticity space involving fold changes (C), plotted in that order for all 224 valid models (see Figure 6G–H). The variance explained by PC1, PC2 and PC3 are mentioned in the respective axes of all PCA plots. The effective dimensionality values (computed as $(\sum \lambda_i)^2 / \sum \lambda_i^2$, for all eigen values $\lambda_i$) computed from PCA was 2.5, 3.95, and 3.95 for the measurements, parametric, and plasticity spaces, respectively.

**Supplementary Table S1: Model parameters with their base-model values.** All channel names and their gating kinetics are from (Beining et al., 2017). The base-model values are from (Kumari and Narayanan, 2024).

| Parameters/Sections | Soma | GCL | IML | MML | OML | AIS | Axon | Count |
| --- | --- | --- | --- | --- | --- | --- | --- | --- |
| **<u>Passive Properties</u>** |  |  |  |  |  |  |  |  |
| ***Intracellular Resistivity, $R_a$ ($\Omega\text{cm}$)*** | | | | | | | | |
| Base-model value | 200 | 200 | 200 | 200 | 200 | 200 | 100 | 2 |
| ***Leak Conductance, $g_{\text{pas}}$ ($\mu\text{S}/\text{cm}^2$)*** | | | | | | | | |
| Base-model value | 13.8 | 13.8 | 20.1 | 26.3 | 26.3 | 6.59 | 6.59 | 4 |
| **<u>Active Properties</u>** |  |  |  |  |  |  |  |  |
| ***Na 8-state channel Maximal Conductance, $g_{\text{Na}}$ ($\text{mS}/\text{cm}^2$)*** | | | | | | | | |
| Base-model value | 18.8 | 2.8 | 2 | 1.25 | 1.25 | 540 | 65.18 | 6 |
| ***Inward rectifier K<sup>+</sup> channel Maximal Conductance, $g_{\text{Kir}}$ ($\text{mS}/\text{cm}^2$)*** | | | | | | | | |
| Base-model value | 0.354 | 0.354 | 0.354 | 0.354 | 0.354 | 0.168 | 0.168 | 2 |
| ***Delayed rectifier K<sup>+</sup> channel Maximal Conductance, $g_{\text{Kv}2.1}$ ($\text{mS}/\text{cm}^2$)*** | | | | | | | | |
| Base-model value | 70.91 | - | - | - | - | - | - | 1 |
| ***Large conductance Ca<sup>2+</sup>-activated potassium (BK) channel properties*** |  |  |  |  |  |  |  |  |
| ***Maximum conductance Alpha subunit, $\alpha_{\text{BK}}$ ($\text{mS}/\text{cm}^2$)*** | | | | | | | | |
| Base-model value | 15.6 | - | - | - | - | 62.4 | 62.4 | 2 |
| ***Maximum conductance Alphabeta subunit, $\alpha\beta_{\text{BK}}$ ($\text{mS}/\text{cm}^2$)*** | | | | | | | | |
| Base-model value | 3.9 | - | - | - | - | 15.6 | 15.6 | 2 |
| ***Small conductance Ca<sup>2+</sup>-dependent potassium (SK) channel Maximal Conductance, $g_{\text{SK}}$ ($\mu\text{S}/\text{cm}^2$)*** | | | | | | | | |
| Base-model value | 0.83 | 1.67 | 4.37 | 4.37 | 4.37 | 83.3 | 12.5 | 5 |
| ***N-type Ca<sup>2+</sup> channel (Cav2.2) Maximal Conductance, $g_{\text{Ca}2.2}$ ($\text{mS}/\text{cm}^2$)*** | | | | | | | | |
| Base-model value | 0.3 | 0.05 | 0.05 | 0.05 | 0.05 | 0.05 | 0.05 | 2 |
| ***Ca<sup>2+</sup> channel (Cav1.2) Maximal Conductance, $g_{\text{Ca}1.2}$ ($\text{mS}/\text{cm}^2$)*** | | | | | | | | |
| Base-model value | 0.02 | 0.01 | 0.04 | 0.04 | 0.04 | 0.01 | - | 3 |
| ***Ca<sup>2+</sup> channel (Cav1.3) Maximal Conductance, $g_{\text{Ca}1.3}$ ($\text{mS}/\text{cm}^2$)*** | | | | | | | | |
| Base-model value | 0.016 | 0.004 | 0.008 | 0.008 | 0.008 | 0.008 | 0.004 | 3 |
| ***T-type Ca<sup>2+</sup> channel (Cav3.2) Maximal Conductance, $g_{\text{Ca}3.2}$ ($\text{mS}/\text{cm}^2$)*** | | | | | | | | |
| Base-model value | 0.022 | 0.022 | 0.022 | 0.022 | 0.022 | 0.008 | 0.008 | 2 |
| ***Ca Buffer - Ca decay constant, $\tau$ (ms)*** | | | | | | | | |
| Base-model value | 240 | 240 | 240 | 240 | 240 | 240 | 43 | 2 |
| ***HCN Channel Maximal Conductance, $g_{\text{HCN}}$ ($\mu\text{S}/\text{cm}^2$)*** | | | | | | | | |
| Base-model value | - | - | 4 | 4 | 4 | - | - | 1 |
| ***K<sup>+</sup> channel (Kv 1.1) Maximal Conductance, $g_{\text{Kv}1.1}$ ($\text{mS}/\text{cm}^2$)*** | | | | | | | | |
| Base-model value | - | - | - | - | - | 0.25 | 0.25 | 1 |
| ***K<sup>+</sup> channel (Kv 1.4) Maximal Conductance, $g_{\text{Kv}1.4}$ ($\text{mS}/\text{cm}^2$)*** | | | | | | | | |
| Base-model value | - | - | - | - | - | 10.12 | 10.12 | 1 |
| ***K<sup>+</sup> channel (Kv 3.4) Maximal Conductance, $g_{\text{Kv}3.4}$ ($\text{mS}/\text{cm}^2$)*** | | | | | | | | |
| Base-model value | - | - | - | - | - | 30.78 | 7.66 | 2 |
| ***A-Type K<sup>+</sup> channel (Kv 4.2) Maximal Conductance, $g_{\text{Kv}4.2}$ ($\text{mS}/\text{cm}^2$)*** | | | | | | | | |
| Base-model value | - | 2.17 | 4.35 | 4.35 | 4.35 | - | - | 2 |
| ***M-type K<sup>+</sup> channel (Kv 7.2/3) Maximal Conductance, $g_{\text{Kv}7.2/3}$ ($\text{mS}/\text{cm}^2$)*** | | | | | | | | |
| Base-model value | - | - | - | - | - | 6.7 | 1.34 | 2 |
| **Total Parameters** |  |  |  |  |  |  |  | **45** |

**Supplementary Table S2: Sub-threshold measurements of base model DG granule cells and their respective electrophysiological bounds.** The bounds were derived from respective electrophysiological measurements reported for dorsal and intermediate granule cells (Mishra and Narayanan, 2020, Kumari and Narayanan, 2026). It may be noted that all base model sub-threshold measurements are within their respective bounds.

|  | Sub-threshold Measurements | Base Model | Lower bound | Upper bound |
| --- | --- | --- | --- | --- |
| 1. | Resting membrane potential, $V_{RMP}$ (mV) | -79.93 | -80 | -70 |
| 2. | Input resistance, $R_{in}$ ($M\Omega$) | 112.34 | 90 | 300 |
| 3. | Maximal impedance amplitude, $\vert Z\vert _{max}$ ($M\Omega$) | 112.81 | 90 | 225 |
| 4. | Resonance frequency, $f_R$ (Hz) | 0.937 | 0.4 | 1.2 |
| 5. | Resonance strength, $Q$ | 1.012 | 1 | 1.2 |
| 6. | Total Inductive Phase, $\Phi_L$ (rad.Hz) | 0 | 0 | 0.03 |
| 7. | Sag (%) | 4.7 | 1 | 7 |
| 8. | Summation ratio of αEPSPs, $S_\alpha$ | 1.06 | 0.9 | 1.5 |

**Supplementary Table S3: Supra-threshold measurements of base model DG granule cells and their respective electrophysiological bounds.** The bounds were derived from respective electrophysiological measurements reported for dorsal and intermediate granule cells (Mishra and Narayanan, 2020, Kumari and Narayanan, 2026). It may be noted that all base model supra-threshold measurements are within their respective bounds.

|  | Supra-threshold Measurements | Base Model | Lower bound | Upper bound |
| --- | --- | --- | --- | --- |
| 1. | Firing frequency at 50 pA, $f_{50}$ (Hz) | 0 | 0 | 0 |
| 2. | Firing frequency at 250 pA, $f_{250}$ (Hz) | 18 | 5 | 35 |
| 3. | Action potential threshold, $V_{th}$ (mV) | -42.92 | -50 | -30 |
| 4. | Action potential amplitude, $V_{AP}$ (mV) | 108.539 | 100 | — |
| 5. | Action potential halfwidth, $T_{APHW}$ (ms) | 1.125 | 0.7 | 1.4 |
| 6. | Peak $dV/dt$, $dV/dt\vert _{max}$ (V/s) | 318.473 | 200 | 700 |
| 7. | Minimum $dV/dt$, $dV/dt\vert _{min}$ (V/s) | -48.335 | -160 | -40 |
| 8. | Latency to first spike, $T_{1AP}$ (ms) | 13.1 | 5 | 100 |
| 9. | First interspike interval, $T_{1ISI}$ (ms) | 48.3 | 5 | 100 |
